# Local externalization of phosphatidylserine mediates developmental synaptic pruning by microglia

**DOI:** 10.1101/2020.04.24.059584

**Authors:** Nicole Scott-Hewitt, Fabio Perrucci, Raffaella Morini, Marco Erreni, Matthew Mahoney, Agata Witkowska, Alanna Carey, Elisa Faggiani, Lisa Theresia Schuetz, Sydney Mason, Matteo Tamborini, Matteo Bizzotto, Lorena Passoni, Fabia Filipello, Reinhard Jahn, Beth Stevens, Michela Matteoli

## Abstract

Neuronal circuits assembly requires the fine equilibrium between synapse formation and elimination. Microglia, through the elimination of supernumerary synapses, have an established role in this process. While the microglial receptor TREM2 and the soluble complement proteins C1q and C3 are recognized key players in this process, the neuronal molecular components that tag synapses to be eliminated are still undefined. Here we show that exposed phosphatidylserine (PS) represents a neuronal ‘eat-me’ signal enabling microglial-mediated synapse pruning. In hippocampal neuron and microglia co-cultures, synapse elimination can be prevented by blocking accessibility of exposed PS using Annexin V or through microglial loss of TREM2. *In vivo*, exposed PS is detectable at both hippocampal and retinogeniculate synapses, where exposure coincides with the onset of synapse elimination and increased PS engulfment by microglia. Mice deficient in C1q, which fail to properly refine retinogeniculate connections, display elevated exposed PS and reduced PS engulfment by microglia. These data provide mechanistic insight into microglial-mediated synapse pruning and identify a novel role of developmentally regulated PS exposure that is common among developing brain structures.

## Introduction

The development of a properly connected nervous system is a complex process involving the regulation of synapse formation, controlled elimination of supernumerary synapses, and maintenance of appropriate connections. Elucidating the mechanisms of synaptic pruning is important to understand normal neurodevelopment and maturation, and how disruptions in this process may contribute to neurological dysfunction. The involvement of microglia, the CNS resident immune cells and phagocytes, in synaptic refinement has been widely established. Microglial processes make direct and transient connections with subsets of neuronal synapses, with evidence for presynaptic and postsynaptic elements inside microglial lysosomes (Basilico *et al*, 2019; Filipello, Morini *et al*, 2018; Paolicelli *et al*, 2011; Schafer *et al*, 2012; Tremblay *et al*, 2010; Weinhard *et al*, 2018). Microglia-mediated synaptic pruning is a highly regulated process and occurs in several brain regions during developmental critical periods of synaptic refinement. In addition to the role in developmental circuit refinement, this process can become activated in vulnerable brain regions in disease models (Hong *et al*, 2016; Neher *et al*, 2012; Schafer *et al*, 2016; Vasek *et al*, 2016; Wang *et al*, 2015), suggesting that disease and development share common regulators and mechanisms of pruning.

Microglia engulf immature synapses and axon terminals through a process involving the microglial complement receptor (CR3/Mac1)(Schafer *et al.*, 2012) and the phagocytic receptor TREM2 (Filipello *et al.*, 2018). However, other synapses and parts of the neuron remain intact, suggesting the existence of local molecular cues that direct the microglia to specific axons or synapses. What are the signals that trigger this process at the appropriate time and place? In the visual system, complement factors C1q, C3, and C4 decorate subsets of synaptic inputs for removal by phagocytic microglia (Schafer *et al.*, 2016; Sekar *et al*, 2016; Stevens *et al*, 2007). However− given that these factors represent secreted soluble proteins− a crucial question is whether neuronal signals exist that mark specific synapses for elimination and whether this mechanism might be shared by different, developing brain regions.

Externalization of phosphatidylserine (PS) has been established as one of the first detectable events to occur in cells undergoing apoptosis. However recent studies have uncovered transient, localized PS exposure events that occur in a non-apoptotic manner (Segawa *et al*, 2011; Smrz *et al*, 2007). For example, in *Drosophila*, it has recently been demonstrated that PS exposure can occur locally on injured dendrites, which are then targeted for elimination while sparing the remaining uninjured cell structures (Sapar *et al*, 2018). The recognition of exposed PS by phagocytes, a process that is critical in the clearance of apoptotic cells and debris to prevent autoimmunity, is facilitated by numerous phagocytic receptors including microglial TREM2 (Graham *et al*, 2014; Grommes *et al*, 2008; Park *et al*, 2007; Shirotani *et al*, 2019). Further, complement binding to exposed PS, either directly or indirectly, can mediate engulfment in the periphery (Martin *et al*, 2012; Païdassi *et al*, 2008). Recent work has shown that exposed PS may be present *ex vivo* on isolated synaptosomes also tagged with C1q (Györffy *et al*, 2018); however whether local externalization of PS occurs *in vivo* in a non-injury associated manner and what role this type of signal would have, is unclear.

We hypothesized that PS exposure occurs locally during normal development, acting as an ‘eat-me’ signal which specifies synapses to be targeted for elimination during microglial-mediated pruning. We found that liposome engulfment by isolated microglia was dependent on PS concentration, and that, when co-cultured with hippocampal neurons, microglial-dependent synaptic reduction also required exposed PS. *In vivo,* synaptic PS occurred predominantly at presynaptic inputs in both the hippocampus and the dorsal lateral geniculate nucleus (dLGN), and was highest during developmental periods coinciding with synapse elimination. In C1q knockout animals we observed an increase in PS exposed at presynaptic inputs as well as a decrease in microglial engulfment of PS-labeled material. Taken together, our data provide mechanistic insights into developmental circuit refinement, identify a novel role of exposed PS in specifying synapses for elimination, and provide a framework for the convergence of several previously described mechanisms of microglial-mediated developmental synaptic pruning.

## Results

### Microglia Phagocytose Liposomes in a Phosphatidylserine-Dependent Manner

We investigated whether the amount of exposed PS correlates with microglia engulfment by taking advantage of uniformly sized synthetic liposomes containing different amounts of PS and cardiolipin, and labeled with the lipid fluorescent dye DiO (shown as percentage, PS_99:1_, PS_50:49:1_; PS_20:79:1_, all numbers are mol%). Liposomes were administered to microglia and their internalization was quantified (Fig. 1A). As expected, liposomes were internalized by CD68-positive phagolysosomal structures (Fig. 1A, B, C, D and E). The extent of PS_99:1_ liposome internalization was significantly lower when microglia were devoid of the phagocytic receptor TREM2 (Fig. 1B, C and F), which recognizes PS (Wang *et al.*, 2015). Of note, microglia displayed a reduced phagocytic activity when the amount of PS in liposomes was lower (phagocytic activity efficiency: PS_99:1_> PS_50:49:1_> PS_20:79:1_) (Fig. 1B, D and G).

**Figure 1:**
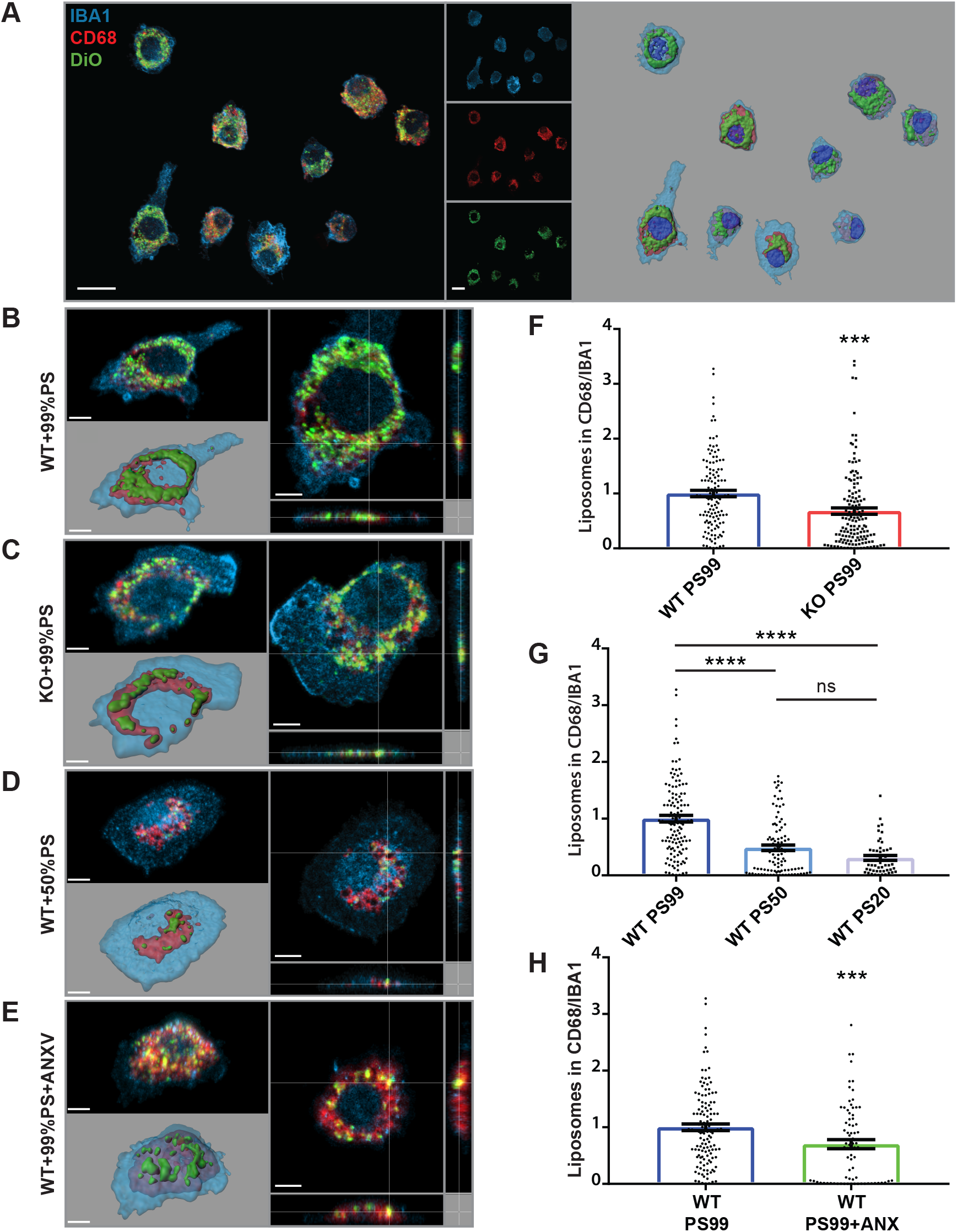
Microglia engulf liposomes in PS-dependent manner. (**A**) Representative confocal images and 3D reconstruction of in vitro WT microglia fed with DiO-labeled liposomes containing a controlled amount of phosphatidylserine (PS; 99%, 50%, 20%). Microglia were stained with Iba1 (blue) and lysosomes labeled with CD68 (red). The amount of DiO-labeled liposomes (green) internalized by microglia (and identified through CD68 co-localization) was quantified using Imaris software. Scale bar: 10 μm. (**B, C, F**) Confocal images showing (B) WT and (C) *Trem2*^−/−^ microglia fed with 99%PS liposomes and (F) relative histogram representing the quantification of the engulfed materials (WT: 1 ± 0.06; KO: 0.68 ± 0.06). (**D, G**) Representative images of (D) WT microglia fed with 50%PS, and (G) related histogram representing quantification of the engulfed materials (PS99: 1 ± 0.06; PS50: 0.49 ± 0.05; PS20: 0.31 ± 0.04). (**E, H**) Representative images of (E) WT microglia pre-treated with ANXV and fed with 99%PS liposomes and (H) related histogram representing quantification of engulfed materials (WT+PS99 w/o ANXV: 1 ± 0.67; WT+PS99+ANXV: 0.70 ± 0.08). Note: KO+PS99 w/o ANX vs KO+PS99+ANXV was not statisically significant (KO+PS99 w/o ANXV: 0.68 ± 0.06 vs KO+PS99+ANXV: 0.57 ± 0.06). Scale bar in (B-E): 3μm. At least three independent experiments were performed; ***p<0.001 ****p<0.0001 One-way ANOVA with Dunn’s multiple comparisons test. Bars in (F-H) represent mean ± SEM.

Cloaking PS by annexin V (ANXV), an innate molecule which binds with high affinity to PS-bearing membranes, significantly reduced engulfment of PS_99:1_ liposomes (Fig 1B, E and H). Conversely, ANXV pre-treatment did not affect the residual phagocytosis of PS_99:1_ liposomes by TREM2-deficient microglia (data in Fig 1 legend). These data indicate that microglia engulf liposomes in a PS-dependent manner.

### Microglial-Mediated Synapse Elimination is Dependent on Exposed PS In Vitro

Microglia eliminate extranumerary synapses in the developing brain, regulating the dynamics of synaptic connections throughout life. It has been previously shown that co-culturing microglial cells in contact with hippocampal neurons for 24 hours results in microglia-mediated synapse elimination (Filipello, Morini *et al.*, 2018), as indicated by the reduction of mushroom spine density (Fig. 2A) and frequency of miniature Excitatory Postsynaptic Currents (mEPSCs) (Fig. 2B). To address whether PS exposure is required for microglia-mediated synapse elimination, hippocampal neurons were exposed to ANXV, 15 minutes before being co-cultured with WT microglia. Cloaking of exposed PS completely prevented both dendritic spine and mEPSC reduction (Fig. 2A and 2B). Neuronal exposure to ANXV did not affect microglia viability, as assessed by calcein-PI assay (Fig. S1A), nor microglial phagocytic activity, as shown by beads phagocytosis assay (Fig. S1B). When microglia from TREM2-deficient mice were cultured with WT neurons, no elimination of synapses was detected, in line with previous results (Filipello, Morini *et al.*, 2018) (Fig. 2C and 2D). In the latter case, cloaking of exposed PS with ANXV did not induce any modification in either spine density or miniature events (Fig. 2C and 2D). As a further control, exposure of pure neuronal cultures to ANXV did not alter either spine density or mEPSC frequency (Fig. S1C). These results indicate that masking exposed PS prevents microglia-mediated synaptic engulfment.

**Figure 2:**
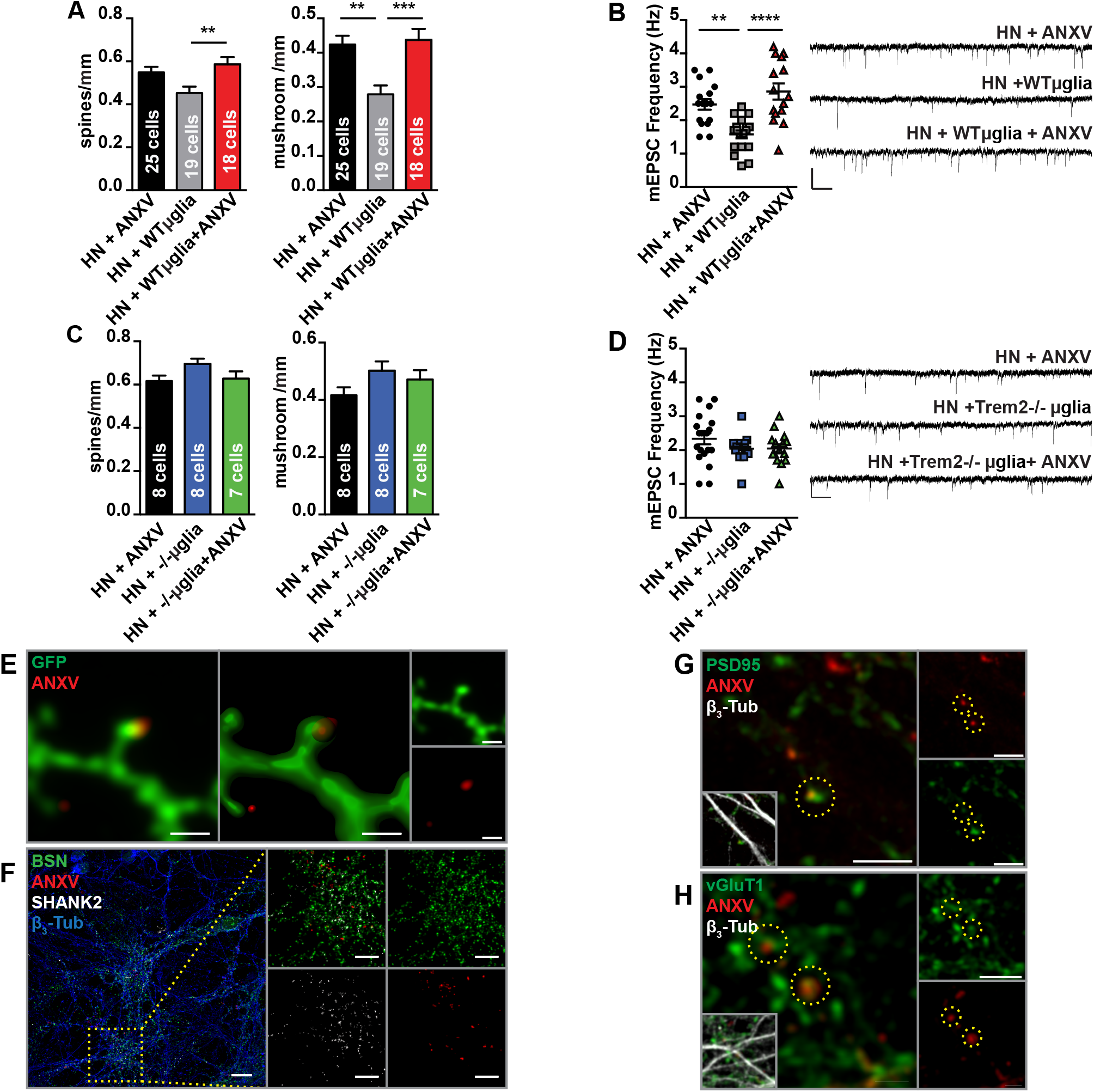
In vitro microglia-mediated synapse elimination is ePS-dependent. **(A)** Quantitative analysis of total and mushroom spines density of hippocampal neurons (HN) alone, co-cultured with WT microglia or exposed to ANXV. Number of total spines per mm, HN+ANXV= 0.6 ± 0.02, number of examined dendrites: 63, number of neurons: 29; HN+WT microglia= 0.52 ± 0.02, number of examined dendrites: 101, number of neurons: 41; HN+WT microglia+ANXV= 0.63 ± 0.16, number of examined dendrites: 94, number of neurons: 38. **(B)** Histogram and representative traces of mEPSCs whole-cell recordings showing HN frequency in the same conditions described for (A). Averaged mEPSC Frequency: HN+ANXV 2.28 ± 0.19 n =16 cells, HN+WT microglia 1.623 ± 0.12, n=16 cells, HN+WT microglia+ANXV 2.86 ± 0.24, n=15 cells. **(C)** Quantitative analysis of total and mushroom spines density of HN alone, co-cultured with Trem2−/− microglia or exposed to ANXV. No statistical differences were found. **(D)** Histogram and representative traces of mEPSCs whole-cell recordings showing HN frequency in the same conditions described for C. No statistical differences were found. Averaged mEPSC Frequency: HN+ANXV: 2.38 ± 0.13, HN+KO microglia 2.2 ± 0.17, n=6 cells, HN+microglia KO+ANXV: 2.03 ± 0.14 n=8 cells. **(E)** GFP expressing neurons labeled with ANXV. Scale bar 1μm. **(F)** Representative confocal images of HN labeled with ANV (red) and stained for the neuronal marker β3-tubulin (blue), the presynaptic marker Bassoon (green) and postsynaptic markers Shank2 (grey). (**G**, **H**) Representative confocal images of β3-tubulin HN (grey) labeled with ANV (red) and stained for PSD95 (green) (G) and vGlut1 (green) (H). Co-localization between ANXV positive puncta and PSD95/ vGlut1 markers is highlighted by yellow dashed circles. No significant differences in ANXV co-localization between pre or postsynaptic markers were detected in HN cultures. scale bar: (E) 10μm; (F) 5μm; (G) 2μm. Note: for (A-D) At least three independent experiments were performed for all condition. ** P < 0.01, ****P<0.0001, One way ANOVA followed by Tukey’s multiple comparison test. Bars represent mean ± SEM. Scale bars, 10 pA and 250ms.

To investigate whether PS is externalized at synaptic sites, primary cultures of hippocampal neurons were exposed to ANXV-568, fixed and stained for the presynaptic markers Bassoon or vGluT1 and the postsynaptic markers Shank2 or PSD95. Confocal microscopy analysis followed by a deconvolution process indicated that ANXV labels a subpopulation of hippocampal synapses in vitro (Fig. 2 F, G and H). To obtain a better visualization of PS exposure, neuronal cultures were transfected with eGFP, which allows the detection of dendritic processes and spines, and subsequently exposed to ANXV. The presence of exposed PS in apposition to dendritic protrusions is shown in Fig. 2E. These data indicate that PS is exposed at synaptic sites *in vitro* and that PS exposure is required for microglial synapse elimination.

### PS is Exposed at Synapses In Vivo in the Developing Hippocampus

To test whether PS exposure occurs locally at synapses *in vivo*, we utilized the commercially available PS-binding probe PSVue (PSVue^®^-550). Once activated by zinc, PSVue has fast, calcium-independent binding kinetics; is significantly smaller in size compared to annexin-based probes, and, similarly to ANXV, is cell impermeable, thus labeling only PS that has been externalized (Smith *et al*, 2011). While PSVue has been used previously to monitor cell death *in vivo* in several disease and injury models (Chan *et al*, 2015; Mazzoni *et al*, 2019; Smith *et al*, 2012), it remains unknown whether localized, non-apoptotic PS exposure occurs *in vivo*.

To assess whether externalized PS is detectable in the developing CA1 hippocampal region, PSVue was intracerebroventricularly (ICV) injected in mice at postnatal day (P)10 and P18, which represent the central time window for synapse refining in hippocampus (Paolicelli *et al.*, 2011), and at P30 (young adult), when refinement is largely completed. Analysis was performed 3 hours following injection (Dallérac *et al*, 2011; Walrave *et al*, 2016). Section preincubation with Sudan Black, which reacts against lipofuscin, an aggregate of oxidized proteins and lipids, reduced autofluorescence (Fig. S2). To directly test whether PS is exposed at synapses in CA1, brain sections were stained for the presynaptic marker Bassoon and the postsynaptic marker PSD-95 (Fig. 3A-B). A subset of synaptic sites in CA1 were labeled by PSVue, with a significantly higher co-localization detected for Bassoon relative to PSD-95 (Fig.3C-D). We examined synaptic PS exposure at several relevant time points throughout this process, specifically P10, P18, and P30. Mice were injected 3 hours prior to each time point with PSVue and analyzed. Results revealed the highest synaptic staining at P10, when hippocampal synapse elimination peaks, with progressively lower labeling at P18 and P30, which is at the end of the pruning period (Fig. 3E-F).

**Figure 3:**
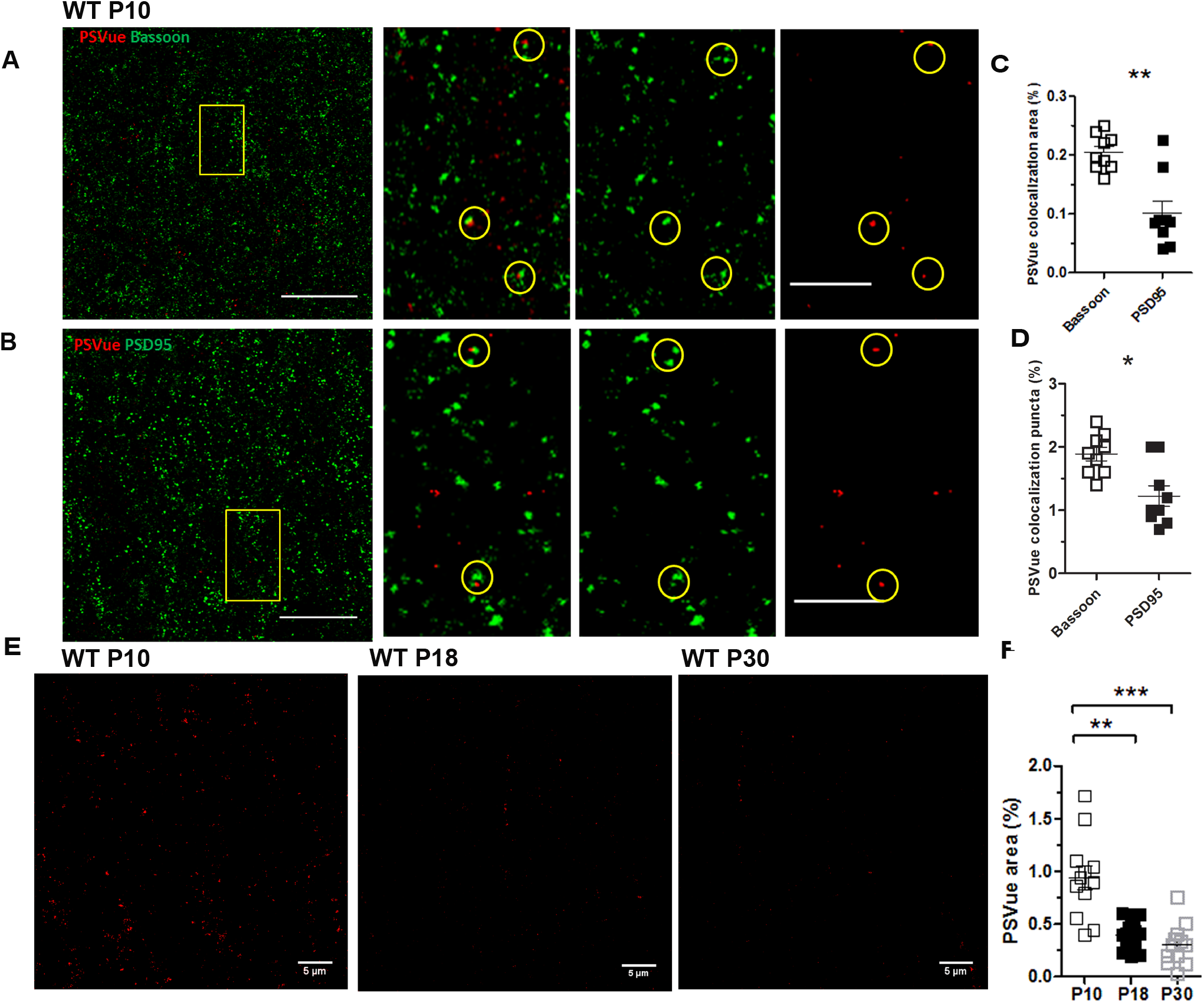
ePS occurs in vivo at synapses. (**A-B**) Representative images of presynaptic (Bassoon) (A) and postsynaptic (PSD95) (B) IHC performed in the CA1 of WT P10 animals sacrificed 3 hours after PSVue injection. Images taken at 63X magnification. Scale bar 5 um. (**C-D**) Quantification of PSVue with either pre- (Bassoon) or postsynaptic (PSD95) markers co-localization in CA1 of WT P10 animals following PSVue injection (3 hours treatment). PSVue+Bassoon colocalized area: 0.2047 ± 0.1 vs PSVue+PSD95 colocalized area: 0.1017 ± 0.02; PSVue+Bassoon colocalized puncta 1.889% ± 0.1 vs PSVue+PSD95 colocalized puncta 1.22% ± 0.16; N=3 animals n=9 fields **p<0.01P10 * p<0,05 **p<0,01 Unpaired t-test **(E)** Representative images of the CA1 hippocampal region of WT P10, P18 and P30 animals sacrificed 3 hours after PSVue injection. Images were acquired with a 63x magnification. Scale bar 5 um. **(F)** Quantification of PSVue signal in the CA1 hippocampal region of WT P10, P18 and P30 animals sacrificed 3 hours after PSVue injection. PSvue area: P10 0.9408% ± 0.11 vs. P18 0.3983% ± 0.03 vs. P30: 0.3017% ± 0.050; **p<0.01, ***p<0.001 P10 N=3 animals n=12 fields; P18 N=4 animals, n=16 fields; P30 N=3 animals n=12 fields. **p<0.01, ***p<0,001 One-way ANOVA with Kruskal-Wallis test.

### Exposed PS is Developmentally Regulated Across Periods of Pruning in the Visual System

To determine whether PS exposure occurred more broadly during periods of developmental pruning throughout the brain, we turned to the visual system, and specifically the dorsal lateral geniculate nucleus (dLGN), where microglial-mediated pruning and synaptic refinement has been extensively studied (Lehrman *et al*, 2018; Schafer *et al.*, 2012; Stevens *et al.*, 2007). We performed ICV injections on C57/Bl6 animals at P4 with PSVue and conducted analyses 24 hours later. Following ICV injection into the left ventricle, PSVue signal was assessed in the contralateral dLGN, where we observed robust, punctate PSVue-labeling. No signal was observed in the dLGN of animals injected with control PSVue non-activated by zinc (Fig. 4A, B).

**Figure 4:**
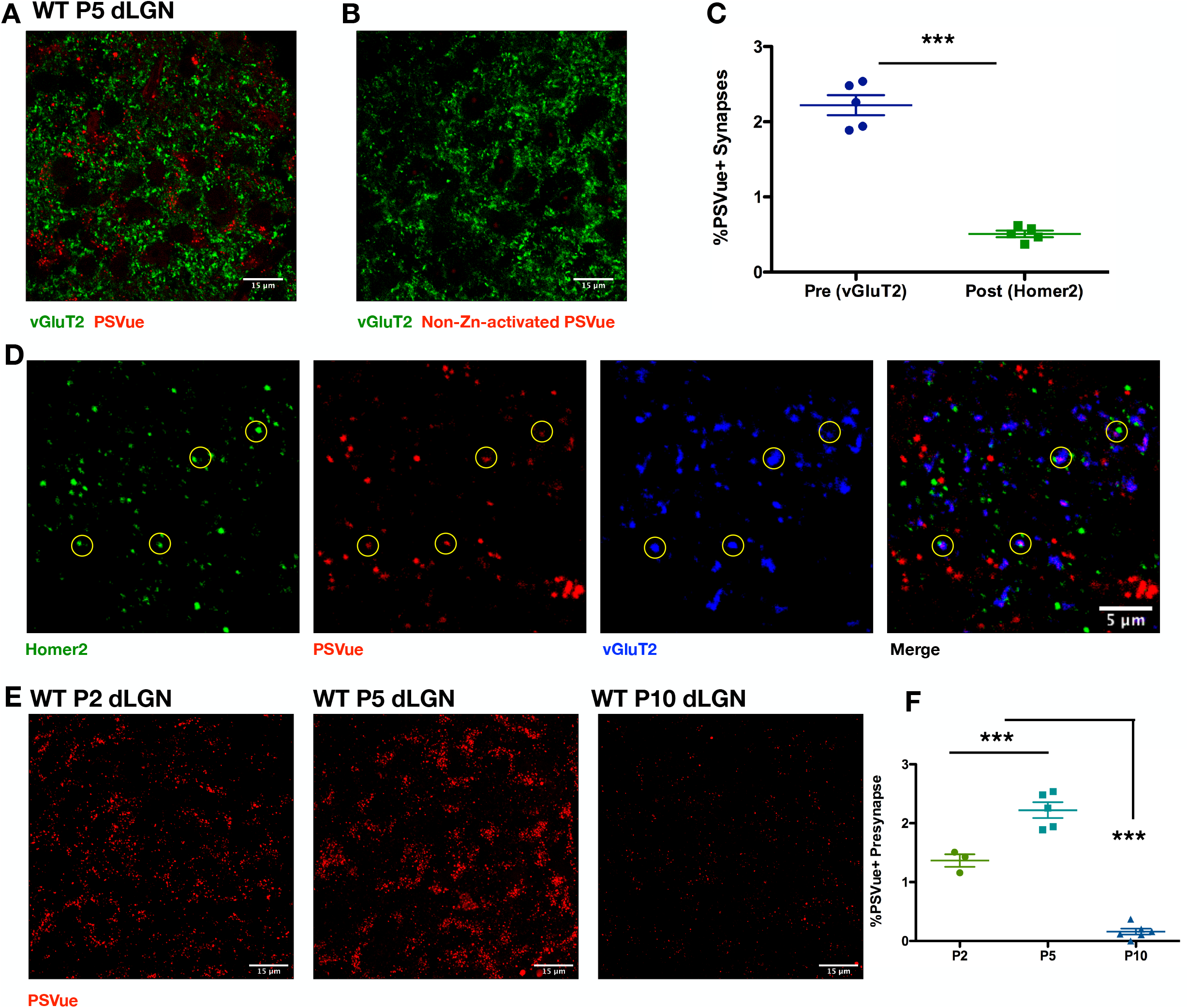
*In vivo* exposed PS is developmentally regulated across periods of pruning in the visual system. **(A, B)** Representative max intensity images of the dLGN following injection with either PSVue (**A**) or non-zinc-activated PSVue (**B**) in WT P4 C57/Bl6 mice 24 hours prior. IHC for the presynaptic marker vGluT2 was performed. Images taken at 63X magnification; scale bar represents 15μm. **(C)** Quantification of PSVue co-localization at synapses in the dLGN of WT P5 mice injected with PSVue 24 hours prior. Data represent the mean per animal ± SEM; N=4; PSVue+Presynaptic: 2.22% ± 0.134 vs. PSVue+Postsynaptic: 0.51% ± 0.043 ***p<0.0001 unpaired t-test. **(D)** Representative images of presynaptic (vGluT2) and postsynaptic (Homer2) IHC performed in the dLGN of WT P5 animals following PSVue injection 24 hours prior. Synapses were identified through co-localization of pre- and postsynaptic markers in CellProfiler. Circles indicate synapses where PSVue co-localization was observed. Images taken at 63X magnification; scale bar represents 5μm. **(E)** Representative max intensity images of the dLGN following injection with PSVue 24 hours prior in WT P2, P5, and P10 C57/Bl6 mice. Images taken at 63X magnification; scale bar represents 15μm. **(F)** Quantification of PSVue co-localization with vGluT2 at synapses in the dLGN of WT P2, P5, and P10 C57/Bl6 mice injected with PSVue 24 hours prior. Data represent the mean per animal ± SEM; N=3 (P2), N=5 (P5), N=6 (P10); P2: 1.36% ± 0.105 vs. P5: 2.22% ± 0.134 vs. P10: 0.16% ± 0.050 ***p<0.0001 One-way ANOVA with Tukey’s Multiple Comparison Test.

To directly test whether exposed PS occurred on synapses in the developing dLGN, we used immunohistochemistry to label presynaptic (vGluT2) and postsynaptic (Homer2) terminals of PSVue-injected animals at P5. By defining synapses as the co-localization of pre- and postsynaptic markers, we identified those that further co-localized with PSVue and, more specifically, determined whether PSVue labeled the presynaptic or postsynaptic component. We found that, similarly to the hippocampus (Fig. 3A, B), PSVue labeled a subset of synapses in the dLGN and was largely localized to presynaptic rather than postsynaptic inputs (Fig. 4B, C).

Finally, we analyzed exposed PS throughout the process of eye segregation in the dLGN, a well-characterized period of developmental pruning where retinal ganglion cell (RGC) inputs from either eye are segregated into discrete patches (Huberman, 2007; Jaubert-Miazza *et al*, 2005). In mice, this occurs between P2−P8 and is largely completed by P10. We examined synaptic PS exposure at several relevant time points throughout this process, specifically P2, P5, and P10. Wild type C57/Bl6 mice were injected 24 hours prior to each time point with PSVue and analyzed. We found robust PSVue labeling at younger ages (P2, P5) that was considerably reduced by P10 (Fig. 4E). Quantification of PSVue presynaptic (vGluT2) co-localization peaked at P5 and was significantly diminished by P10 (Fig. 4F). Taken together, our data demonstrate that PS is exposed *in vivo* at synapses, coinciding with periods of developmental synaptic refinement in several regions.

### In Vivo Developmental PS Exposure is not Caspase 3-Dependent

Programmed RGC apoptosis occurs early postnatally, typically between P0−P2, in a caspase 3-dependent manner (Cellerino *et al*, 2000). To ensure that the exposed PS observed *in vivo* in the dLGN was not due to cells undergoing apoptosis, we performed staining for activated (cleaved) caspase 3 in PSVue-injected animals. No signal for active caspase 3 was observed in either the dLGN or the retina (Fig. S3A, B; respectively). To further examine whether developmental PS exposure was downstream of caspase 3 activation, we performed PSVue ICV injections, as described above, in P4 WT and caspase 3 KO littermates (Jax #006233). When analyzed 24 hours later at P5, no significant difference in PSVue labeling was observed in the dLGN between WT and caspase 3 KO littermates (Fig. S3C, D), revealing that PS exposure observed *in vivo* is not downstream of caspase 3-mediated activation or apoptosis.

### Microglia Engulf PSVue-Labeled Inputs In Vivo in the dLGN and Hippocampus

The role of microglia in synaptic pruning has been established in several developing brain regions, including the visual system (both the dLGN (Lehrman *et al.*, 2018; Schafer *et al.*, 2012; Stevens *et al.*, 2007) and primary visual cortex (Tremblay *et al.*, 2010)) and the hippocampus (Filipello *et al.*, 2018; Paolicelli *et al.*, 2011; Weinhard *et al.*, 2018). Given that we observed that i) exposed PS was necessary for microglial-mediated synaptic reductions *in vitro* (Fig. 1, 2), and ii) PSVue labeling occurs *in vivo* across periods of developmental pruning at synapses in both the hippocampus (Fig. 3) and dLGN (Fig. 4), we hypothesized that exposed PS may broadly act as a synaptic ‘eat-me’ signal, leading to microglial recognition and elimination.

To test whether microglia engulf PSVue-labeled material in the dLGN and hippocampus, we utilized our established method of *in vivo* engulfment analysis (Schafer *et al*, 2014) by measuring the volume of PSVue co-localized with the microglial lysosomal marker CD68 within microglia. Wild type C57/Bl6 mice were injected 24 hours prior to each time point with PSVue and analyzed by immunohistochemistry. We observed that microglia (labeled by Iba1 and P2y12 antibodies) within the dLGN (Fig. 5A, B) and hippocampus (Fig. 5D, E) had internalized PSVue-labeled material. Microglial engulfment was temporally regulated in both regions. In the dLGN, engulfment of PSVue-labeled material was highest at P5 and was significantly reduced by P10 (Fig. 5C), similar to what was observed previously when quantifying microglial engulfment of presynaptic inputs in the dLGN (Schafer *et al.*, 2012). In the hippocampus, we evaluated engulfment at P10 and P30, time points where microglial pruning has been observed (P10) and completed (P30) (Filipello, Morini *et al.*, 2018; Paolicelli *et al.*, 2011), in both CA1 (Fig. 3C, D) and CA3 (Fig. S4A-C) regions. Similar to the dLGN, engulfment was significantly higher in the hippocampus of P10 animals when compared to P30 (CA1 Fig. 5F; CA3 Fig. S4D−F). Taken together, these data reveal that microglia engulf PSVue-labeled material *in vivo* in a temporally regulated manner during development, coinciding with established periods of microglial synaptic pruning.

**Figure 5:**
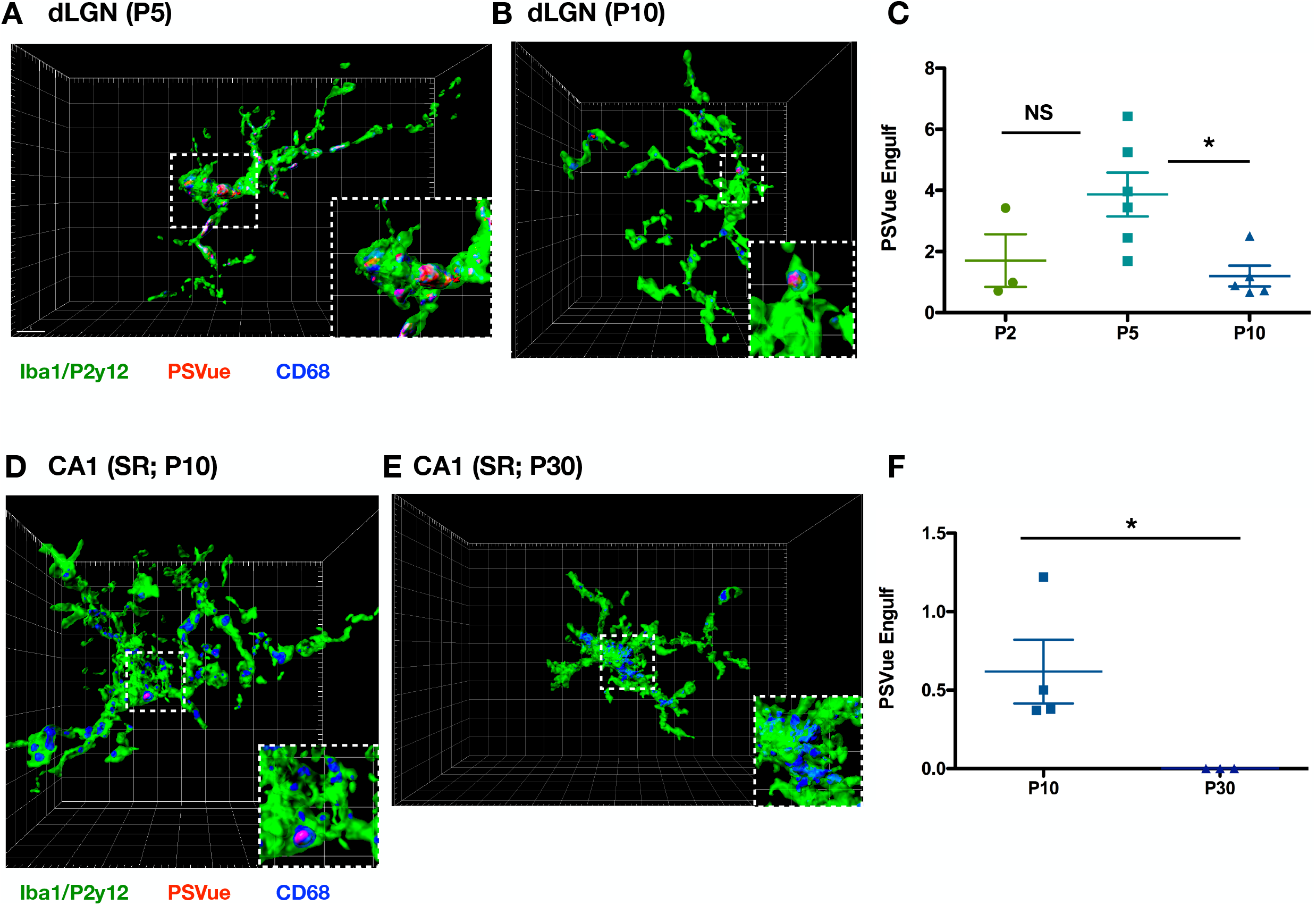
Microglia engulf PSVue-labeled inputs *in vivo* in the dLGN and hippocampus. **(A, B)** Representative Imaris surface-rendered images of microglia in the dLGN following injection with PSVue 24 hours prior in WT P5 **(A)** and P10 (**B)** C57/Bl6 mice. Microglia are labeled by IHC with Iba1 and P2y12, and lysosomes are labeled with CD68. Images taken at 63X magnification. **(C)** Quantification of the volume of engulfed PSVue material in dLGN microglia analyzed from WT P2, P5, and P10 C57/Bl6 mice injected with PSVue 24 hours prior. Data represent the mean of 15-20 microglia per animal ± SEM; N=3 (P2), N=5 (P5), N=6 (P10); P2: 1.703% ± 0.861 vs. P5: 3.868% ± 0.715 vs. P10: 1.196% ± 0.340 *p=0.0237 (P5 vs. P10) One-way ANOVA with Tukey’s Multiple Comparison Test. **(D, E)** Representative Imaris surface-rendered images of microglia in the SR of CA1 following injection with PSVue 24 hours prior in WT P10 **(D)** and P30 **(E)** C57/Bl6 mice. Microglia are labeled by IHC with Iba1 and P2y12, and lysosomes are labeled with CD68. Images taken at 63X magnification. **(F)** Quantification of the volume of engulfed PSVue material in CA1 microglia analyzed from WT P10 and P30 C57/Bl6 mice injected with PSVue 24 hours prior. Data represent the mean of 15-20 microglia per animal ± SEM; N=4 (P10), N=3 (P30); P10: 0.617% ± 0.203 vs. P30: 1.17e^−5^% ± 3.33e^−7^ *p=0.05 One-way ANOVA with Tukey’s Multiple Comparison Test.

### Increased PSVue Labeling and Microglial Engulfment of Ipsilateral RGC Inputs During Eye Segregation

As eye segregation proceeds in the dLGN, contralateral and ipsilateral RGC inputs are segregated into two distinct regions. Although inputs from both eyes are removed, the area occupied by ipsilateral inputs is more greatly reduced (Jaubert-Miazza *et al.*, 2005). To test whether PS exposure occurred preferentially on ipsilateral inputs we performed intraocular injections with fluorescently labeled CTB, labeling contralateral (CTB-488) and ipsilateral (CTB-647) RGC inputs of P4 WT C57/Bl6 mice. Concurrent PSVue ICV injections were performed, and the dLGN was analyzed 24 hours later at P5 (Fig. 6A). As reported (Jaubert-Miazza *et al.*, 2005), we found that the percentage of dLGN volume occupied by contralateral RGC inputs was significantly higher relative to ipsilateral RGC inputs (Fig. 6B). However, when we calculated the percentage of PSVue-labeled CTB inputs, we found that a significantly higher percentage of ipsilateral inputs (CTB647) were co-localized with PSVue (Fig. 6C).

**Figure 6:**
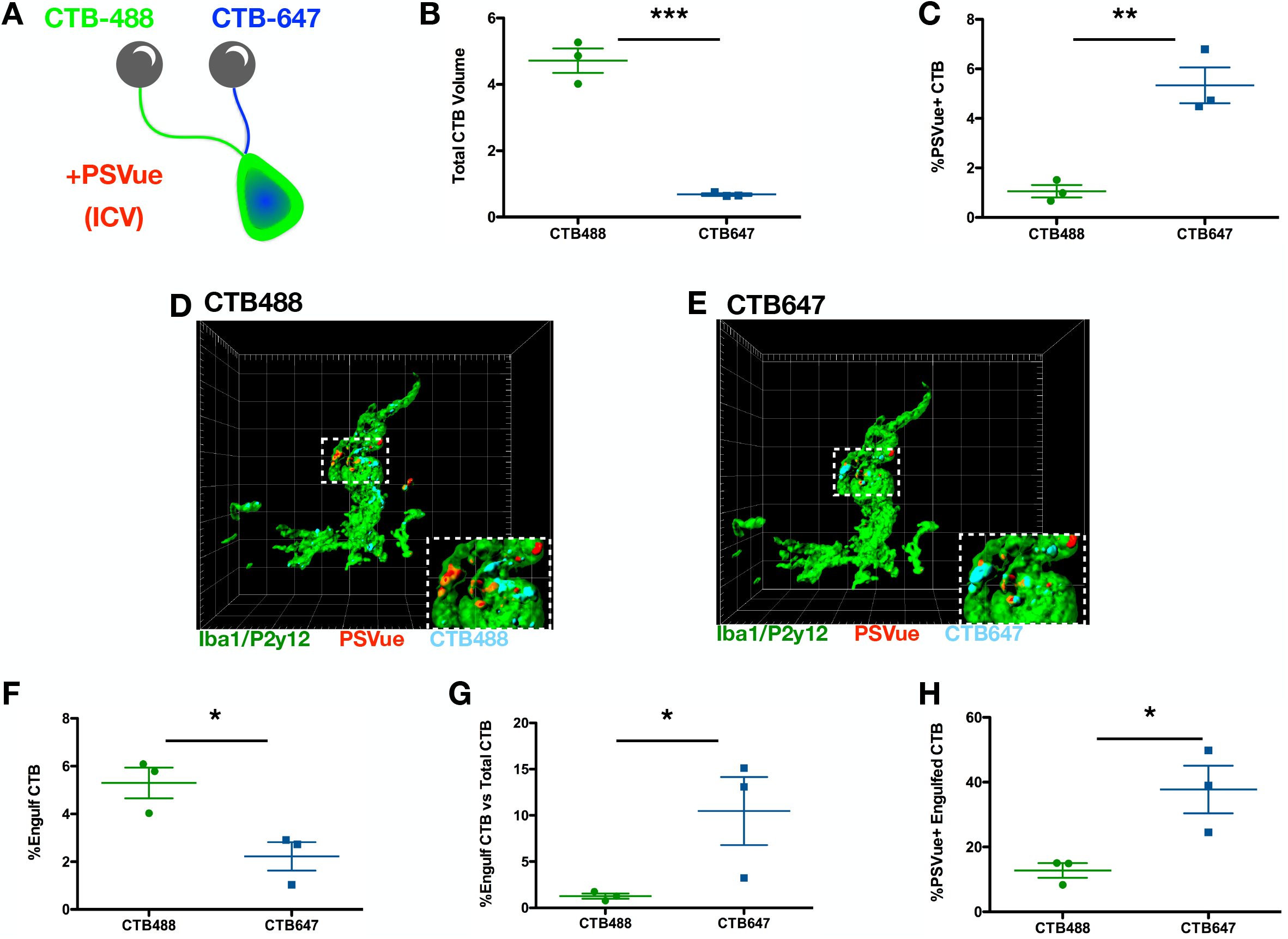
Increased PSVue labeling and microglial engulfment of ipsilateral RGC inputs during eye segregation. **(A)** Schematic of the experimental design for analysis of PSVue co-localization onto retinal ganglion cell (RGC) inputs. Cholera toxin β subunit (CTB) conjugated to Alexa 488 or 647 was intraocularly injected into the contralateral or ipsilateral eyes of P4 WT mice, respectively. PSVue was then ICV injected and animals were sacrificed 24 hours later. **(B)** Quantification of the total volume of CTB-488 or CTB-647 RGC inputs in the dLGN of P5 mice injected with PSVue 24 hours prior. Data represent mean volume per animal. N=3 animals ± SEM; CTB-488: 4.717% ± 0.367 vs. CTB-647: 0.68% ± 0.035 ***p=0.0004 unpaired t-test. **(C)** Quantification of the percentage of the volume of CTB-488 or CTB-647 RGC inputs that co-localized with PSVue. The volume of co-localized signal was divided by the total volume of the inputs for either eye. N= 3 animals ± SEM; CTB-488: 1.06% ± 0.247 vs. CTB-647: 5.337% ± 0.73 **p=0.0052 unpaired t-test. **(D, E)** Representative Imaris surface-rendered images of a microglia showing engulfed PSVue material along with engulfed CTB-488 **(D)** or CTB-647 **(E)** in the dLGN following injection with PSVue 24 hours prior in WT P5 mice. Microglia are labeled by IHC with Iba1 and P2y12. Images taken at 63X magnification. **(F)** Quantification of the volume of engulfed CTB-488 or CTB-647 material in dLGN microglia analyzed from WT P5 mice injected 24 hours prior. Data represent the mean engulfment of either CTB-488 or CTB-647 from 15-20 microglia per animal ± SEM; N=3, CTB-488: 5.3% ± 0.641 vs. CTB-647: 2.22% ± 0.597 *p=0.0246 unpaired t-test. **(G)** Quantification of the volume of microglial engulfed CTB-488 or CTB-647 normalized to the total volume of contralateral or ipsilateral CTB. Data represent normalized CTB engulfment of CTB-488 orCTB-647 from 15-20 microglia per animal ± SEM; N=3, CTB-488: 1.27% ± 0.280 vs. CTB-647: 11.33% ± 2.83 *p=0.0241 unpaired t-test. **(H)** Quantification of the volume of co-localized PSVue and CTB (488 vs 647) engulfed by microglia normalized to the volume of engulfed CTB-488 or CTB-647. Data represent normalized PSVue+/CTB+ engulfment of contralateral (488) vs ipsilateral (647) from 15-20 microglia per animal ± SEM; N=3, CTB-488: 12.77% ± 2.231 vs. CTB-647: 37.76% ± 07.331 *p=0.0311 unpaired t-test.

To determine whether elevated ipsilateral PS exposure translated into preferential microglial elimination, we directly compared PSVue-labeled contralateral and ipsilateral engulfment within dLGN microglia (Fig. 6D, E). We found that microglia engulfed significantly more contralateral inputs (CTB-488) when compared to ipsilateral (CTB-647), when the volume of internalized CTB was normalized to microglia volume (Fig. 6F). As described above, the total volume of ipsilateral inputs was significantly less than the volume of contralateral inputs. When we took this into account by normalizing the volume of engulfed CTB by the total CTB volume, we found that a higher proportion of ipsilateral inputs within the dLGN had been engulfed (Fig. 6G). Furthermore, when we calculated the percentage of engulfed CTB that also co-localized with PSVue, we found that significantly more engulfed ipsilateral inputs were positive for PSVue (Fig. 6H). These data reveal that, although ipsilateral inputs account for far less volume of total RGC inputs in the developing dLGN, exposed PS occurs on significantly more of them, resulting in their disproportionate elimination by microglia.

### Loss of C1q Leads to Elevated PSVue-Positive Inputs and Reduced Microglial PSVue Engulfment

Microglia engulf presynaptic RGC inputs within the developing dLGN in a complement-dependent manner (Schafer *et al.*, 2012; Stevens *et al.*, 2007). The soluble complement proteins C1q and downstream C3 label a subset of retinogeniculate synapses in the dLGN, targeting them for elimination. How these proteins bind to specific synapses remains unknown. However, in the periphery, complement proteins have been found to bind to PS (either directly (Païdassi *et al.*, 2008) or indirectly (Martin *et al.*, 2012)) to facilitate phagocytic removal. Given our findings that local synaptic PS exposure occurs during periods of developmental pruning, we hypothesized that exposed PS is the signal that leads to synaptic C1q deposition and subsequent microglial elimination. Immunohistochemistry analysis of P5 WT C57/Bl6 mice injected with PSVue 24 hours prior revealed that C1q and PSVue co-localized at a subset of retinogeniculate synapses (Fig. 7A), and analysis across several developmental time points revealed that co-localization of PSVue with C1q at presynaptic inputs (vGluT2) peaked at P5 (Fig. 7B). Interestingly, we also observed considerable overlap between C1q and PSVue that was not co-localized to vGluT2 (Fig. 7A). Thus, the role of non-synaptic C1q, and how local exposed PS may interact with it during development, remains an area for further investigation.

**Figure 7:**
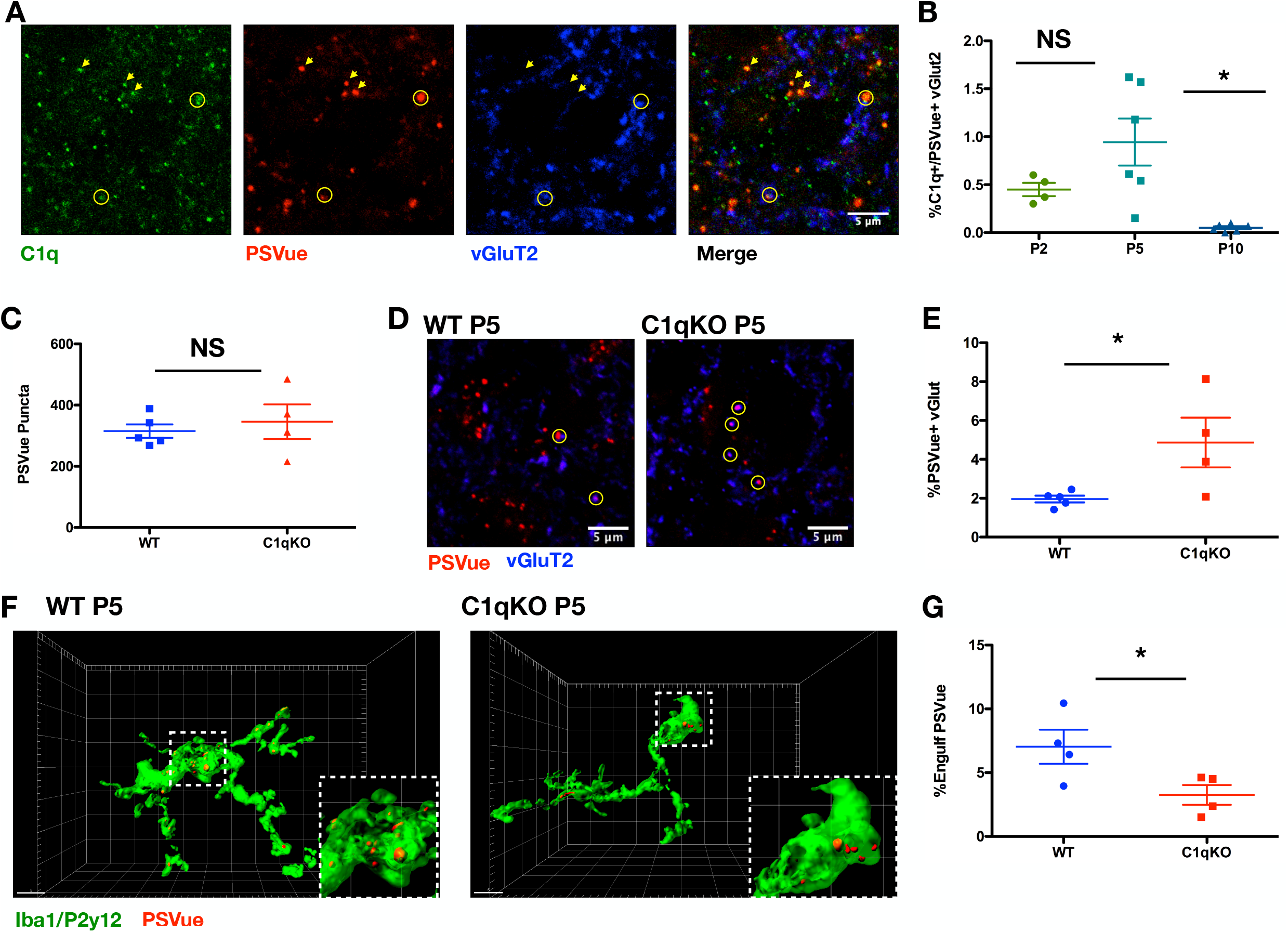
Loss of C1q leads to increased PSVue+ inputs and reduced microglial PSVue engulfment. **(A)** Representative IHC images of C1q and vGluT2 in the dLGN of WT P5 animals following PSVue injection 24 hours prior. Circles indicate where PSVue co-localization with C1q and vGluT2 was observed. Arrows indicate where PSVue and C1q co-localization was observed independent of vGluT2. Images taken at 63X magnification; scale bar represents 5μm. **(B)** Quantification of PSVue co-localization with vGluT2 and C1q in the dLGN of WT P2, P5, and P10 C57/Bl6 mice injected with PSVue 24 hours prior. Data were calculated by quantifying the percentage of PSVue+/C1q+/vGluT2+ puncta normalized to total vGluT2. Data represent the mean per animal ± SEM; N=4 (P2), N=6 (P5), N=5 (P10); P2: 0.45% ± 0.069 vs. P5: 0.945% ±0.245 vs. P10: 0.051% ± 0.015 *p=0.047(P5 vs. P10) One-way ANOVA with Tukey’s Multiple Comparison Test. **(C)** Quantification of PSVue in the dLGN of WT or C1qKO P5 littermates injected with PSVue 24 hours prior. Data represent the mean per animal ± SEM; N=5 (WT), N=4 (C1qKO); WT: 315.0 ± 22.04 vs. C1qKO: 345.8 ± 56.47 p=0.597 unpaired t-test. **(D)** Representative images of the dLGN of P5 WT or C1qKO littermates following injection with PSVue 24 hours prior. IHC was performed for the presynaptic marker vGluT2. Circles indicate where co-localization between PSVue and vGluT2 was observed. Images taken at 63X magnification; scale bar represents 5μm. **(E)** Quantification of PSVue co-localization with vGluT2 at synapses of P5 WT or C1qKO littermates following injection with PSVue 24 hours prior. Data represent the mean per animal ± SEM; N=5 (WT), N=4 (C1qKO); WT: 1.962% ± 0.174 vs. C1qKO: 4.864% ± 1.277 *p=0.0382 t-test. **(F)** Representative Imaris surface-rendered images of microglia in the dLGN following injection with PSVue 24 hours prior in P5 WT or C1qKO littermates. Microglia are labeled by IHC with Iba1 and P2y12. Images taken at 63X magnification. **(G)** Quantification of the volume of engulfed PSVue material in microglia of P5 WT or C1qKO littermates following injection with PSVue 24 hours prior. Data represent the mean of 15-20 microglia per animal ± SEM; N=4 (WT), N=4 (C1qKO); WT: 6.978% ± 1.294 vs. C1qKO: 3.258% ± 0.775 *p=0.0487 unpaired t-test.

Mice lacking C1q have disrupted developmental pruning, resulting in impaired eye segregation and excessive retinal innervation of the dLGN (Stevens *et al.*, 2007). To test whether PS exposure and microglia engulfment were also affected in these animals, we performed PSVue ICV injections into P4 WT and C1qKO littermates and analyzed 24 hours later. We found that, although the total amount of PSVue was unchanged between WT and C1qKO (Fig. 7C), a significantly higher percentage of vGluT2 presynaptic inputs were labeled with PSVue in C1qKO animals (Fig. 7D, E). Finally, analysis of P5 WT and C1qKO littermates revealed significantly reduced microglial PSVue engulfment in animals deficient in C1qKO (Fig. 7F, G). All together, these data demonstrate that C1q deficiency leads to elevated presynaptic PSVue and reduced microglial engulfment, and suggest that synaptic PS exposure may act as the signal leading to C1q deposition during pruning.

## Discussion

Microglia eliminate synapses during developmental pruning, but the neuronal signals specifying which synapses are targeted remain unknown. Here we demonstrate that local synaptic PS exposure enables microglial recognition and elimination. We found that liposome engulfment by isolated microglia was dependent on PS concentration and TREM2 *in vitro*. In co-cultures with hippocampal neurons, microglial-dependent synaptic reduction was also PS- and TREM2-dependent. *In vivo,* we utilized the PS-binding probe PSVue and found that PS exposure occurred locally at synapses in both the hippocampus and the dLGN early in development and that microglia in both regions engulfed PSVue-labeled material, a process that was also developmentally regulated. Neuronal PS exposure was not stochastic, as ipsilateral inputs in the dLGN were preferentially labeled and targeted for engulfment by microglia. Finally, in the dLGN, presynaptic C1q and PSVue co-localization peaked during developmental microglial pruning, and in C1q knockout animals we observed an increase in PSVue-labeled presynaptic inputs as well as a decrease in microglial engulfment of PSVue-labeled material. Our findings integrate known mechanisms of microglial-mediated synaptic refinement, namely the microglial receptor TREM2 (Filipello, Morini *et al.*, 2018) and the complement pathway (Schafer *et al.*, 2012; Sekar *et al.*, 2016; Stevens *et al.*, 2007), by identifying local PS exposure as a common neuronal signal triggering engulfment across multiple developing brain regions. In addition to further elucidating the molecular underpinnings of microglial-mediated synaptic pruning, our data provide mechanistic insight into how these pathways may contribute to neurological disorders.

### Emerging Roles of PS Localization and Regulation in Diverse Biological Functions

PS is a ubiquitous lipid, estimated to represent 13-15% of all phospholipids in the human cerebral cortex (Svennerholm, 1968). Along with other anionic lipids, PS is typically asymmetrically distributed within the inner leaflet where several critical functions have been ascribed. PS participates in membrane translocation and activation of protein kinase C (Newton & Keranen, 1994), is involved in Akt signalling through interactions with PIP3 (Huang *et al*, 2011), and contributes to membrane curvature and vesicular transport (Xu *et al*, 2013). At the synaptic level, PS alters the interaction of SNARE proteins with membranes (Dennison *et al*, 2006), influences neurotransmitter vesicle fusion (Murray *et al*, 2004), binds to synaptotagmins (Fernandez *et al*, 2001; Zhang *et al*, 2009) and modulates AMPA glutamate receptors (Baudry *et al*, 1991).

While the function of inner leaflet plasma membrane PS continues to be an area of active research, emerging roles of PS that has been exposed on the outer leaflet are also being discovered. PS exposure is classically associated with apoptosis, where loss of membrane asymmetry occurs early on in the cell-death signaling cascade (Fadok *et al*, 1992; Hoffmann *et al*, 2001; Martin *et al*, 1995). Non-apoptotic PS exposure has also been described in several biological contexts. In the blood, PS exposure occurs on activated platelets, where it contributes to the clotting response following injury (Lentz, 2003). In the immune system, transient PS exposure occurs during the activation of T lymphocytes, B lymphocytes, mast cells and neutrophils (Callahan *et al*, 2003; Elliott *et al*, 2005; Martin *et al*, 2000). In neurons, PS exposure can occur locally on injured axons (Shacham-Silverberg *et al*, 2018) or dendrites (Sapar *et al.*, 2018) which are then targeted for elimination while sparing the remaining uninjured cell structures. Ectopic PS exposure on living neurons causes phagocytic cells to engulf distal neurites (Sapar *et al.*, 2018), and direct masking of exposed PS signal reduces axonal debris engulfment (Shacham-Silverberg *et al.*, 2018). Our data reveal an additional novel role of locally externalized PS, where synaptic exposure contributes to microglial-mediated synaptic pruning, a critical process important for the maturation of neuronal circuits during development.

An array of enzymes, namely flippases and scramblases, control PS membrane asymmetry and localization; the upstream mechanisms regulating these proteins, and whether apoptotic and local PS exposure are similarly controlled, remain largely unknown. Flippases, also known as type-IV P-type ATPases (P4-ATPases), catalyze the movement of anionic phospholipid species from the extracellular leaflet to the cytosolic leaflet. Alternatively, scramblases function to disrupt membrane asymmetry in a calcium-dependent, ATP-independent manner. Many different flippases and scramblases have been identified to-date, with much of the current understanding of upstream regulation focused on caspase-mediated inactivation and activation, respectively. For example, PS is irreversibly exposed on apoptotic cells by the activation of a phosphatidylserine scramblase, Xk-related protein 8 (Xkr8), through caspase cleavage (Suzuki *et al*, 2016) and inhibition of phosphatidylserine translocases ATP11A and ATP11C is also caspase-dependent (Segawa *et al*, 2014). While caspase activation appears to be an important mechanism in apoptosis-associated PS exposure, it is likely not the upstream mechanism driving the local synaptic exposed PS that we observed *in vivo*, as revealed by our studies in caspase 3 deficient mice (Fig. S3).

A critical question that remains is the identification of the upstream mechanism driving synaptic PS exposure during development. Previous work has demonstrated that in some cells transient PS exposure relies on TMEM16 family phosphatidylserine scramblases and/or calcium inhibition of translocases ATP11A or ATP11C (Suzuki *et al*, 2010). Interestingly, Anoctamin 1 (ANO1)/TMEM16A, a Ca2^+^-activated Cl^−^ channel, has been shown to mediate process extension in RGCs, contributing to cortex development (Hong *et al*, 2019). Whether the TMEM16 family of scramblases could be involved in PS exposure at synapses is worth investigating.

The role of neuronal activity has been well established in synaptic refinement (Burbridge *et al*, 2014; Hooks & Chen, 2006; Hua & Smith, 2004; Huberman, 2007) and microglial-mediated synaptic pruning (Schafer *et al.*, 2012), but how activity is translated into local cues mediating engulfment, including synaptic PS exposure, is not well understood. In the dLGN, although both contralateral and ipsilateral RGC inputs are pruned by microglia during the process of eye segregation, the area occupied by ipsilateral inputs is more significantly reduced (Jaubert-Miazza *et al.*, 2005). Interestingly, we found that ipsilateral RGC inputs were preferentially labeled with PSVue, and these inputs were disproportionately targeted for microglial elimination. Of note, in both the hippocampus and dLGN, exposed PS signal was significantly higher during the first two postnatal weeks, revealing that the upstream mechanisms leading to synaptic PS exposure show specificity and are developmentally regulated. We recently demonstrated that CD47, a well-known ‘don’t eat me’ signal also involved in microglial pruning, was regulated in an activity-driven manner (Lehrman *et al.*, 2018); how activity contributes to synaptic PS exposure and concomitant loss of CD47, and whether these signals are intrinsically linked, is an area of research we are currently testing.

### Mechanisms of Glial Synaptic Pruning Converge on PS as a Common Cue

Numerous phagocytic receptors involved in the recognition of exposed PS (either directly or through adapter proteins such as Gas6) are expressed by microglia and astrocytes, including TREM2, MerTK, Axl, Tyro3, and Bai1 (Chung *et al*, 2013; Fadok *et al*, 2001; Graham *et al.*, 2014; Grommes *et al.*, 2008; Nakano *et al*, 1997; Neher *et al.*, 2012; Park *et al.*, 2007; Païdassi *et al.*, 2008; Shirotani *et al.*, 2019). Further, several of these receptors have been implicated in synaptic elimination by glia (Chung *et al.*, 2013; Filipello, Morini *et al.*, 2018). For example, TREM2 mediates synaptic pruning in the developing hippocampus; where loss of TREM2 resulted in compromised brain connectivity in adult mice (Filipello, Morini *et al.*, 2018). TREM2, for which PS is a known ligand (Wang *et al.*, 2015), was also necessary for PS recognition and internalization (Fig.1). Additionally, components of the complement pathway mediate microglial-mediated synaptic pruning of retinal inputs (Schafer *et al.*, 2012; Stevens *et al.*, 2007; Hong et al., 2016). Here we show that C1q and exposed PS labeled a subset of retinogeniculate synapses *in vivo*, and that C1q-deficient mice had elevated presynaptic exposed PS and reduced levels of PSVue microglial engulfment (Fig. 7D-G). Together our findings suggest that exposed PS could be a common signal linking several pathways involved in glial-mediated synaptic pruning; whether and how these pathways cooperate throughout this process across multiple developing brain regions, and whether astrocytes also contribute to PS removal, will be interesting to examine further.

### Potential Role of Exposed PS and Microglial Engulfment in Neurological Diseases

Emerging evidence suggests developmental mechanisms of synaptic pruning can become aberrantly ‘reactivated’ and contribute to pathological synapse loss and dysfunction (Salter & Stevens, 2017; Stephan *et al*, 2012). In Alzheimer’s Disease (AD), for example, both the complement cascade and TREM2 have been implicated in pathology (Dejanovic *et al*, 2018; Hong *et al.*, 2016; Parhizkar *et al*, 2019; Wang *et al.*, 2015). Indeed, synaptic C1q is aberrantly elevated in several models of AD, contributing to early synapse loss through microglial-mediated elimination (Dejanovic *et al.*, 2018; Hong *et al.*, 2016). In synaptosomes isolated *ex vivo* from AD patient tissue, elevated PS exposure has been reported (Bader Lange *et al*, 2008); whether this also occurs *in vivo*, facilitating C1q deposition, remains unknown. On the other hand, GWAS analyses have identified in AD patients several TREM2 variants, most associated with either reduced surface expression or reduced phagocytic activity (Reviewed in (Ulrich *et al*, 2017)). Further, studies in TREM2 deficient AD mouse models have revealed that the receptor is important for the microglial response to amyloid pathology and that impairment of TREM2 function accelerates early amyloidogenesis due to reduced phagocytic clearance of amyloid seeds (Parhizkar *et al.*, 2019). As PS exposure is hypothesized to occur on plaques, TREM2 may have dual roles in AD, where activation through exposed PS binding likely contributes to plaque containment and clearance, while synaptic PS exposure may contribute to aberrant synapse loss. Our findings identify a role of synaptic PS exposure in both C1q- and TREM2-mediated microglial elimination. Understanding the potential interactions between these two mechanisms in development and disease will be an important area of investigation in the context of many neurological diseases, including Alzheimer’s.

## Materials and Methods

### Mouse Strains

#### Humanitas

Mice were housed in the SPF animal facility of Humanitas Clinical and Research Center in individually ventilated cages. All experiments followed the guidelines established by the European Directive 2010/63/EU and the Italian D.Lg. 26/2014. The study was approved by the Italian Ministry of Health. All efforts were made to minimize the number of subjects used and their suffering. C57BL/6 Trem2^−/−^ mice, generated as previously described (Turnbull *et al*, 2006), were provided by Bioxell-Cosmo Pharmaceutical (Milan, Italy) (Correale *et al*, 2013). P10, P18 and P30 male and female littermate animals were used for each experiment. Trem2^−/−^ and littermate control WT mice (backcrossed 12 generations to the C57BL/6 background) were obtained from Marco Colonna (Turnbull *et al.*, 2006) and the two strains were bred in parallel.

#### Boston Children’s Hospital

All mice were used at the ages specified in the experimental procedures outlined below and, unless stated otherwise, male and female mice were included in all experiments. Wildtype C57BL/6 mice were obtained from Charles River, unless otherwise specified below. Caspase 3^−/−^ mice were obtained from Jackson Labs (Jax #006233) and Caspase 3^+/−^ *x* Caspase 3^+/−^ crosses were used to generate wildtype and Caspase 3^−/−^ littermates. C1qa^−/−^ mice are on a C57BL/6J background and were a kind gift from M. Botto (Imperial College, London). C1qa^+/−^ *x* C1qa^+/−^ crosses were used to generate wildtype and C1q^−/−^ littermates. Animals were group-housed in Optimice cages and maintained in the temperature range and environmental conditions recommended by AAALAC. All procedures were approved by the Boston Children’s Hospital institutional animal care and use committee in accordance with NIH guidelines for the humane treatment of animals.

### Neuronal Cultures

Mouse hippocampal neurons were established from hippocampi of embryonic stage E17 fetal mice. Neurons were plated onto glass coverslips coated with poly-L-lysine at densities of 95 cells/mm2. Cells were maintained in Neurobasal medium (Invitrogen) with B27 supplement and antibiotics, 2 mM glutamine and glutamate. All experiments were performed at 13-15 days in vitro (DIV). ANXV cloaking was performed for 15 minutes at a dilution of 1:20.

### Microglial Cultures

Primary microglia were obtained from mixed cultures established from telencephalon of P1-4 mice. Cells were maintained in DMEM containing 20% of heat-inactivated fetal bovine serum (FBS), glucose 0.6%, sodium-pyruvate 2mM and antibiotics. Microglia were isolated at DIV10-15 by shaking flasks at 245 rpm for 45 minutes, then cells were seeded on poly-L-ornithine pre-coated slides of 18mm diameter. For the microglia-neuron co-culture experiments, WT or Trem2^−/−^ microglia were added to hippocampal neurons (13-14 DIV) at a microglia to neuron ratio of 1.5:1 for 24h. In a distinct set of experiments, ANXV blocking treatment was performed by pre-incubating hippocampal neurons with ANXV (1:20) for 15’. After ANXV wash, microglia were added to neurons for 24h. To visualize neuronal processes and dendritic spines, hippocampal neurons were transfected with GFP using Lipofectamine 2000 (Invitrogen) at DIV11.

For the experiments of bead phagocytosis, microglia was incubated with ANXV (1:20) for 15’ and fluorescent particles (3um) (Spherotech) were added to microglial cells at a 1:1 microglia/beads ratio for 2hr. Cells were then washed with ice-cold PBS, fixed and stained with IBA1 antibody and analyzed. To assess microglial viability following treatment with annexin V, microglia were incubated with ANXV (1:20) for 15’, then washed with PBS and tested for viability after 24h using the live marker Calcein combined with propidium iodide (PI) to label dying cells.

### Phagocytosis Assay of Liposomes

Liposomes were prepared by mixing chloroform stocks of cardiolipin and phosphatidylserine (PS) (both 18:1, Avanti Polar Lipids), supplemented with 1mol% DiO (Thermo Fisher Scientific) in given molar ratios (79:20:1; 49:50:1; 0:99:1) followed by drying of the lipid film with a N_2_stream and desiccation under 15—25 mbar for more than 1 h. Lipid film was then rehydrated with a buffer containing HEPES 20 mM (pH 7.4) and KCl 150 mM with agitation on a thermomixer heated to 72°C for at least 1 h. This vesicle mixture was then extruded 25× through polycarbonate membranes with pore size of 400 nm and 100 nm, consecutively. Uniform liposome size among different lipid compositions was confirmed with dynamic light scattering (DynaPro Titan, Wyatt Technology). If needed, blocking of liposomes with ANXV was performed at room temperature for 15 minutes before cell incubation. Microglia were seeded in DMEM containing 5% of heat-inactivated FBS, and after 1h it was lowered to 1%. Liposomes were then added to medium in a final lipid concentration of 0.1mM and after 1h of incubation, cells were fixed and stained for IBA1 and CD68 in order to perform confocal microscopy.

### Immunocytochemistry and Cell Imaging

Cells were fixed for 10 min at room temperature (RT) in 4% (w/v) PFA, 4% (w/v) sucrose, 20 mM NaOH and 5 mM MgCl2 in PBS, pH 7.4. For microglia intracellular staining, cells were permeabilized and blocked for 45 min at RT in 15% (w/v) goat serum, 0.3% (v/v) Saponin, 450 mM NaCl, 20 mM phosphate buffer, pH 7.4 and incubated at 4°C overnight with primary antibodies Iba-1 (1:750, Wako); CD68 (1:1000 Biolegend) diluted in blocking buffer. Following, cells were incubated with secondary antibodies (1:400, Alexa Fluor®-conjugated, Invitrogen) at room temperature for 2h, counterstained with DAPI and mounted in Fluorsave (Calbiochem). For neuronal staining, if needed, cells were stained by treating with ANXV (1/80) for 7 minutes before being fixed, thereafter they were permeabilized and blocked in 15% (w/v) goat serum, 0.3% (v/v) Triton-X100, 450 mM NaCl, 20 mM phosphate buffer, pH 7.4 and stained for SHANK2 (1:1000), VGlut1 (1:2000), beta-tubulin (1:80), PSD-95 (1:500). Confocal images were acquired with a laser scanning LEICA SP8 STED3X confocal microscope, using a HC PL APO 100X/1.40 oil white objective (Leica). Emission filter bandwidths and sequential scanning acquisition were set up, in order to avoid any possible spectral overlap between fluorophores. Images shown in Fig. 2 were deconvolved using Huygens Professional software (SVI, Scientific Volume Imaging) and subsequently processed using Imaris software (Bitplane).

### In Vitro Liposome Engulfment Quantification

Liposomes, CD68 and Iba-1 volumes were quantified by applying 3D surface rendering of confocal stacks in their respective channels, using identical settings (fix thresholds of intensity and voxel) within each experiment. For quantification of liposome engulfment by microglia, only liposome signals present within microglial CD68+ structures were considered, in order to guarantee that only liposomes completely phagocytosed by microglia were included in the analysis. To this aim, a new channel for ‘‘CD68-liposomes’’ was created, by using the mask function in Imaris, masking the DiO signal within CD68+ structures. Quantification of volumes for ‘engulfed liposome’ was performed following the ‘3D Surface rendering of engulfed material’ protocol, previously published by (Schafer *et al.*, 2014). To account for variations in cell size, the amount of ‘engulfed liposome’ was normalized to the volume of the microglia (given by Iba-1+ volume). Total cytosolic liposome volume per cell from the same confocal stacks was also quantified following the same protocol. All data were normalized to WT+PS_99_% values within each experiment.

### Electrophysiology; Cell Culture Recordings

Whole cell voltage-clamp recordings were performed on WT embryonic hippocampal neurons maintained in culture for 13-15 DIV. During recordings, cells were bathed in a standard external solution containing (in mM): 125 NaCl, 5 KCl, 1.2 MgSO4, 1.2 KH2PO4, 2 CaCl2, 6 glucose, and 25 HEPES-NaOH, pH 7.4. Recording pipettes (resistances of 3-5 MU) were filled with a standard intracellular solution containing (in mM): 135 K+-gluconate, 1 EGTA, 10 HEPES, 2 MgCl2, 4 MgATP, and 0.3 Tris-GTP, (pH 7.4). For mEPSC recordings, 1mM tetrodotoxin, 20mM Bicuculline and 50mM AP5 were added to the standard extracellular solution to block spontaneous action potential propagation by GABA-A and NMDA receptors. Recordings were performed in voltage clamp mode at a holding potential of −70 mV using a Multiclamp 700B amplifier and pClamp-10 software (Axon Instruments, Foster City, CA). Series resistance ranged from 10 to 20 MU and was monitored for consistency during recordings. Cells in culture with leak currents >100 pA were excluded from the analysis. Signals were amplified, sampled at 10 kHz, filtered to 2 or 3 KHz, and analyzed using the pClamp 10 data acquisition and analysis program.

### Intracerebroventricular PSVue Injection

For all injections, a volume of 2μL of 1mM PSVue^®^-550 (PSVue; P-1005 Molecular Targeting Technologies; prepared following manufacturer’s instructions) was utilized. No gross toxic effects, overt signs of distress or pain, or any other behavioral changes were observed as expected with ICV injection procedures.

#### Neonates (P0-P9)

Neonatal pups (P0-P5) were anesthetized by hypothermia or isoflurane (P9; 2-4% exposure followed by 1.5% throughout the experiment). Once anesthetized, as determined by lack of toe pinch reflex, PSVue was injected at a location approximately 0.7 - 1.0 mm lateral to the sagittal suture and 0.7 - 1.0 mm caudal from the neonatal bregma with a 27G needle inserted 2mm into the tissue perpendicular to the skull surface. PSVue was injected slowly and the needle was removed 10 – 20 sec after discontinuing the plunger movement to prevent backflow. Following injection, neonates were warmed on a warm water circulating heating pad, and once pink and breathing were returned to the dam. Animals were sacrificed 24 hours later for analysis as described below.

#### P10/P18/P30 (hippocampal synaptic analysis)

Mice were anesthetized using a ketamine (100mg/kg)/xylazine (20mg/kg) cocktail. PSVue or vehicle (Zinc Nitrate solution) with Fluorophore only were stereotaxically (KOPF instruments) injected into left lateral ventricle of two distinct group of animals via a Hamilton syringe at 0.5 μl/min (Gauge33 Hamilton cat# 549-0700). The stereotaxic coordinates were the following: P10: 0.4 mm anteroposterior (AP); 1 mm mediolateral (ML); 1.8 mm dorsoventral (DV) from bregma; P18/P30: 0.5 mm AP, 1.1 mm ML and 2.5 mm DV from bregma. Upon completion of injection (< 15 min), mice were allowed to recover from anesthesia and returned to their home cage. Animals were sacrificed 3 hours later for analysis as described below.

#### P30 (microglial engulfment analysis)

Once anesthetized using isoflurane (2-4% exposure followed by 1.5% throughout the experiment), PSVue was stereotaxically injected into left lateral ventricles (−0.4 mm anteroposterior, 1 mm mediolateral and −2.5 mm dorsoventral) via a Hamilton syringe at 0.5 μl/min similar to ICV procedures described previously (Hong *et al.*, 2016). Upon completion of injection (< 15 min), mice were allowed to wake up from anesthesia and returned to their home cage. Animals were sacrificed 24 hours later for analysis as described below.

### Immunohistochemistry

#### Hippocampal Immunohistochemistry

Mice were anesthetized with a ketamine (100mg/kg)/xylazine (20 mg/kg1) cocktail and perfused transcardially with 0.9% saline, brains were collected and post-fixed in 4% paraformaldehyde for 24h, then washed and transferred to PBS solution for 24-48h at 4°C. 50mm -thick coronal sections were cut at the level of the dorsal hippocampus with a VT1000S Vibratome. Immunofluorescent staining was carried out on free-floating sections. Sections were processed by 45 min RT blocking in 10% Normal Goat serum, 0.2% Triton X-100 and incubated overnight with specific primary antibodies VGLUT1 (1:1000, SySy, Cat# 135 304); Bassoon (1:400, SySy) followed by incubation with secondary antibodies (1:400, Alexa Fluor-conjugated, Invitrogen), counterstained with DAPI and mounted in Fluorsave (Calbiochem). For PSD-95 (1:300, Thermofisher, Cat# 516900) staining, slices were permeabilized at RT in 0.5% Triton X-100 (Sigma), followed by 1 hr RT blocking in 2% BSA 0.5% Triton X-100 and overnight incubation with primary antibody.

Sections were treated for 7 mins with Sudan Black at a concentration of 0.1% in 70% ethanol and then thoroughly washed to reduce tissue autofluorescence and background, while preserving the specific fluorescence signal

#### dLGN Immunohistochemistry

Mice were anesthetized with Avertin (240 mg/kg, i.p.) and transcardially perfused with PBS. Brains and eyes were harvested and post-fixed in 4% PFA (Electron Microscopy Sciences, catalog #15710) for 2h, then washed and transferred to 30% sucrose solution for 24-48h at 4°C. Cryosections were prepared from tissue embedded in a 2:1 mixture of 30% sucrose: OCT (Sakura Finetek, catalog #4583) onto X-tra slides (Leica Microsystems, catalog #3800200). Tissue sections were dried, washed in PBS, and blocked for 2h at room temperature with slow agitation in 20% normal goat serum (Sigma-Aldrich, catalog #G9023–10ML) with 0.3% TX-100 (Sigma-Aldrich, catalog #T8787–100ML) in PBS. Primary antibody (see below) was applied overnight, in 10% normal goat serum with 0.3% TX-100 in PBS, at 4°C with slow agitation. Tissue sections were then washed 3× 10 min with PBS and incubated with the appropriate AlexaFluor-conjugated secondary antibody (1:200; Invitrogen/ThermoFisher Scientific) for 2h, in 10% normal goat serum with 0.3% TX-100 in PBS, at room temperature with slow agitation. Finally, tissue sections were mounted with VECTASHIELD with DAPI (Vector Laboratories, catalog #H-1000). Images were acquired on a LSM880 confocal microscope (Zeiss) at 63X magnification. At least two z-stack images were taken (0.3μm slices) per section and two sections per animal were analyzed.

#### Synaptic labeling

Brains were sectioned at 15μm on a cryostat, and primary antibodies (guinea pig anti-vGluT2, 1:1000 Millipore #AB2251; rabbit anti-Homer2, 1:500 Cedarlane Labs #160203(Sy); mouse anti-Bassoon, 1:300 Enzo Life Sciences # ADI-VAM-PS003-F) for synaptic staining was performed as described above.

#### Active caspase3

Brains were sectioned at 15μm on a cryostat and primary antibody labeling (rabbit anti-cleaved caspase3 (Asp175) 1:500 Cell Signaling Technology #9661S) was performed as described above.

#### Microglia

Brains were sectioned at 40μm on a cryostat, and primary antibodies (rabbit anti-Iba1, 1:500 Wako #016-26461; rabbit anti-P2y12, 1:500 Anaspec #AS-55043A; rat anti-CD68, 1:200 Serotec #MCA1957) for microglial labeling were performed as described above.

#### C1q synaptic labeling

Brains were sectioned at 15μm on a cryostat, and primary antibodies (guinea pig anti-vGluT2, 1:1000 Millipore #AB2251; rabbit anti-C1q, 1:500 Abcam #ab182451) for synaptic staining was performed as described above.

### Retinal Whole Mounts

Whole mount retina preparations were performed by dissecting out the retinas of PSVue-injected mice following transcardial-perfusion. Retinas were blocked in staining buffer (10% Normal Goat Serum and 2% Triton X-100 in PBS) for 1 hour before incubation with the pan-RGC marker Rbpms (guinea pig anti-Rbpms 1:500 HuaBio #HM0401) and active caspase3 (rabbit anti-cleaved caspase3 (Asp175) 1:500 Cell Signaling Technology #9661S) at 4°C for three days. After 3× 10 min washes in PBS, retinas were incubated in appropriate AlexaFluor-conjugated secondary antibodies (1:200; Invitrogen/ThermoFisher Scientific) at room temperature for 1h. Retinas were washed 3× 10 min and mounted using glycerol.

### Image Analysis

#### dLGN Analysis

Confocal images were processed using the open-source programs Ilastik (Berg *et al*, 2019; v1.3.2) and CellProfiler (McQuin *et al*, 2018; v3.0). Ilastik is machine-learning software for image processing that can be trained to recognize various features in a semi-automated manner. Single-channel images were segmented following training in Ilastik, generating binary probability images. These images were then further processed in CellProfiler, cell image analysis software that we utilized to identify objects and generate co-localization data. Specific analyses are described below.

#### PSVue Quantification

PSVue puncta quantification of segmented probability images was performed using CellProfiler, where objects between 3 and 100 pixels were enumerated for each image. The mean per image (two z-stacks per section, two sections per animal) of PSVue puncta was calculated.

#### PSVue synaptic co-localization

To determine PSVue co-localization on synapses, binary probability images were generated for each independent immunostaining in Ilastik and further processed in CellProfiler. PSVue objects were then masked onto either presynaptic (vGluT2) or postsynaptic (Homer2) images. Co-localized objects were then further masked onto postsynaptic or presynaptic images, respectively, to both identify synapses (co-localization of pre- and postsynaptic markers) and determine specific PSVue localization on synapses. The percentage of pre- or postsynaptic PSVue co-localization was calculated by dividing the number of triple positive masked objects by the number of co-localized pre- and postsynaptic objects. The mean percentage per image (two z-stacks per section, two sections per animal) was then calculated.

#### C1q and PSVue presynaptic co-localization

To determine C1q and PSVue co-localization with presynaptic marker vGluT2, binary probability images were generated for each independent immunostaining in Ilastik and further processed in CellProfiler. PSVue objects were masked onto C1q objects, and co-localized objects were then further masked onto presynaptic images. The percentage of presynaptic C1q and PSVue co-localization was calculated for each image (two z-stacks per section, two sections per animal) by dividing the number of triple positive objects by the number of vGluT2 objects.

#### Hippocampal Analysis

Confocal images were acquired with the laser scanning confocal microscope LEICA SP8, using a HC PL APO 63x/NA 1.40 oil objective. Acquisition parameters (i.e., laser power, gain and offset) were kept constant among different conditions in each single experiment. Images were acquired in stratum radiatum of the CA1 subfield of the hippocampus, as indicated. Each image consisted of a stack of images taken through the z-plane (0.5μm). Confocal images were processed and analyzed using ImageJ program and Fiji (NIH software).

#### PSVue 550 quantification

confocal images were modified as 8-bit images and processed applying fixed color balance and threshold mean value by using the *image adjust* function of ImageJ. The resulting images were analyzed by choosing the *measure or analyze particle* function of ImageJ or Fiji. Area fraction is the percentage of pixels in the image or selection that have been highlighted in red using *the image adjust threshold* function divided by the total number of pixels in the image.

#### Co-localization analysis

Co-localization of PSvue with presynaptic marker Bassoon or postsynaptic marker PSD95 as indicated in Fig 3 was measured using the boolean function ‘and’ for the selected channels by choosing the image calculator process function of ImageJ. The resulting image was binarized and used as a co-localization mask to be subtracted from a single channel. Area fraction and/or the number of puncta resulting from co-localization mask subtraction were measured for each marker (Menna *et al*, 2013). PSVue/Bassoon co-localization or PSVue/PSD-95 co-localization puncta were calculated divided by the number of co-localization puncta by the total number of Bassoon and PSD-95 puncta respectively.

### Engulfment Analysis

#### PSVue Engulfment

Mice were injected with PSVue (as described above) and their brains were harvested 24h later using the same methods as for immunohistochemistry. Brains were cryosectioned and 40μm sections were immunostained for microglial markers Iba-1 and P2y12, as well as for the microglial lysosomal marker CD68 (as described above). For each animal, two sections were imaged and used for analysis. Images were acquired on a LSM880 confocal microscope at 63X with 0.3μm z-steps. For engulfment analysis in the dLGN, at least 4 cells were imaged in the ipsilateral territory and at least 4 cells were imaged in the contralateral territory (minimum 8 cells per dLGN, 16-20 cells per animal). Images were processed and analyzed as described previously using the 3D surface rendering software Imaris (Schafer *et al.*, 2014). Data were used to calculate percent engulfment (volume engulfed PSVue/volume of the cell). All experiments were performed blind to genotype when appropriate.

#### CTB and PSVue Engulfment

P4 mice were injected with anterograde tracers (CTB-488 and CTB-594, ThermoFisher Scientific (C-22841, C-22842)) in the left and right eyes, respectively. PSVue ICV injections were also performed as described above, and their brains were harvested 24h later at P5. Brains were sectioned, stained, and imaged as described above. For each microglia quantified, the percent engulfment of CTB-488 and CTB-647 alone and co-localized with PSVue was calculated, along with the input volumes (total volume CTB inputs/volume of field of view).

### Statistical Analysis

PRISM (Graphpad Software) was used to perform Student’s unpaired t-test and analysis of variance (ANOVA), as appropriate. In order to control for multiple comparisons, we used Tukey’s Multiple Comparison Test and Dunn’s multiple comparisons test. Specific statistical analyses can further be found in figure legends.

## Acknowledgements

We thank Yvanka de Soysa and Kevin Mastro for critical comments on the manuscript and Elisabetta Menna (IN-CNR), Davide Pozzi (Humanitas University), and Chandana Kondapalli (BCH) for discussion. We thank Sivapratha Nagappan Chettiar for technical assistance in stereotactic ICV injections of adult animals; Cassandra-Victoria Innocent of the Cellular Imaging Core at Boston Children’s Hospital (C. Chen; NIH U54 HD090255) for technical support with imaging; Milanka Stevanovic for contributions to microglial engulfment quantifications; Arnaud Frouin and Joel Cuadrado for help with mouse colony maintenance (BCH). This work was supported by NIH T32-AG000222 (N.S.H.), NIH R01-NS071008 (B.S), NIH Conte Center (B.S.), PRIN (Ministero dell’Istruzione dell’Università e della Ricerca, #2017A9MK4R) and MinSal FR 2016 (Ministero della Salute #RF-201602361571) to M.M and CARIPLO grant 2018 (#2018-0364) to F.F. F.P. was supported by Fondazione Umberto Veronesi ‘2017 grant’, M.T is supported by Fondazione Veronesi Fellowship 2019.

## Author Contributions

N.S.H, F.P., B.S., and M.M. designed the study and wrote the manuscript, with help from all authors. N.S.H, F.P. and R.M. performed most experiments and data analysis; F. F. contributed to experimental design and analysis; M.E. performed confocal analysis; A.W. and R.J. designed and executed the liposome preparation; L.T.S. performed liposomes treatments and imaging analysis; M.M., S.M., and M.B. performed immunohistochemistry and microglial engulfment analysis, E.F. and M.T. performed stereotactic ICV injections; L.T.S. and M.B. performed hippocampal immunocytochemistry; A.C. performed retinal whole mounts and immunohistochemistry.

