## Supplemental Figure 1 for "Local externalization of phosphatidylserine mediates developmental synaptic pruning by microglia"

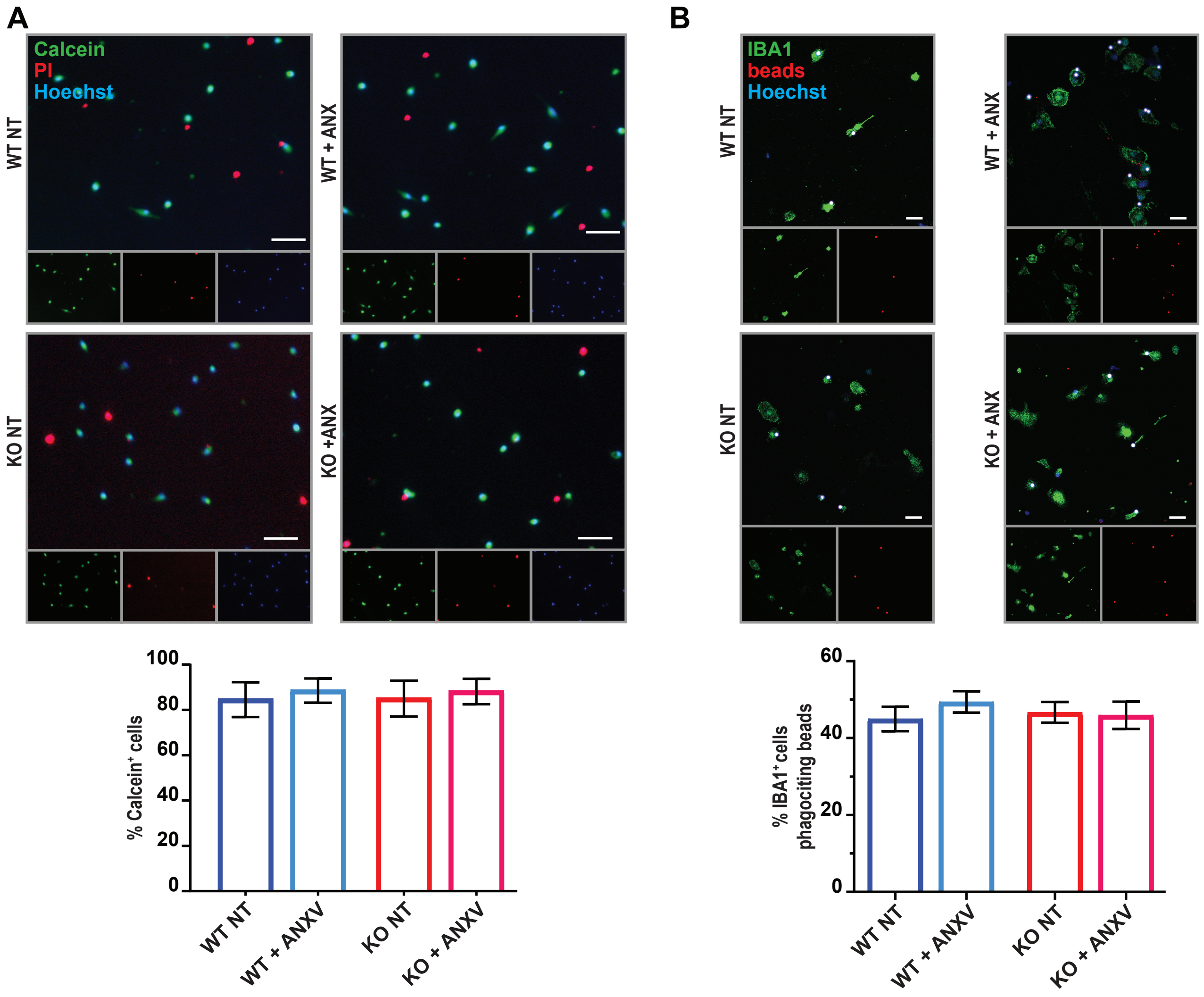


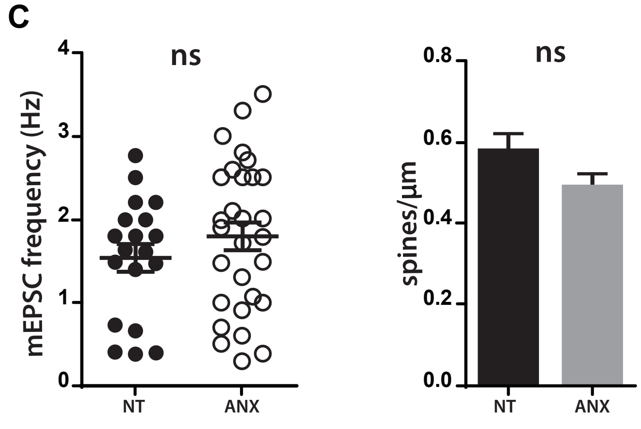


**Supplemental Figure 1: In vitro treatment with Annexin V does not alter microglial viability or properties**.

(**A**) In vitro analysis of microglial viability following PS-binding protein Annexin V (ANXV) treatment. Viability was determined using the live marker Calcein (gree) combined with propidium iodide (PI) (red) to label dying cells. No significant difference was observed between untreated and treated microglia (WT NT: 84.54% ± 7.67; WT+ANXV: 88.5% ± 5.36; KO NT: 84.94% ± 7.93; KO+ANXV: 88.12 ± 5.63). Three independent experiments were performed; scale bar 30µm.

(**B**) In vitro analysis of microglial phagocytic activity following ANXV treatment. Phagocytosis was assessed by measuring microglial engulfment of fluorescent beads (3µm; Spherotech). No significant difference was observed between untreated and treated microglia (WT NT: 44.95 ± 3.19; WT+ANXV: 49.39 ± 2.76; KO NT: 46.68 ± 2.73; KO+ANXV: 45.93 ± 3.54). Three independent experiments were performed; scale bar 10µm.

(**C**) In vitro analysis of mEPSC frequency and spine density of hippocampal neurons following ANXV treatment. No significant difference was observed between untreated and treated neurons for either parameter. Three independent experiments were performed.
