## Supplemental Figure 2 for "Local externalization of phosphatidylserine mediates developmental synaptic pruning by microglia"

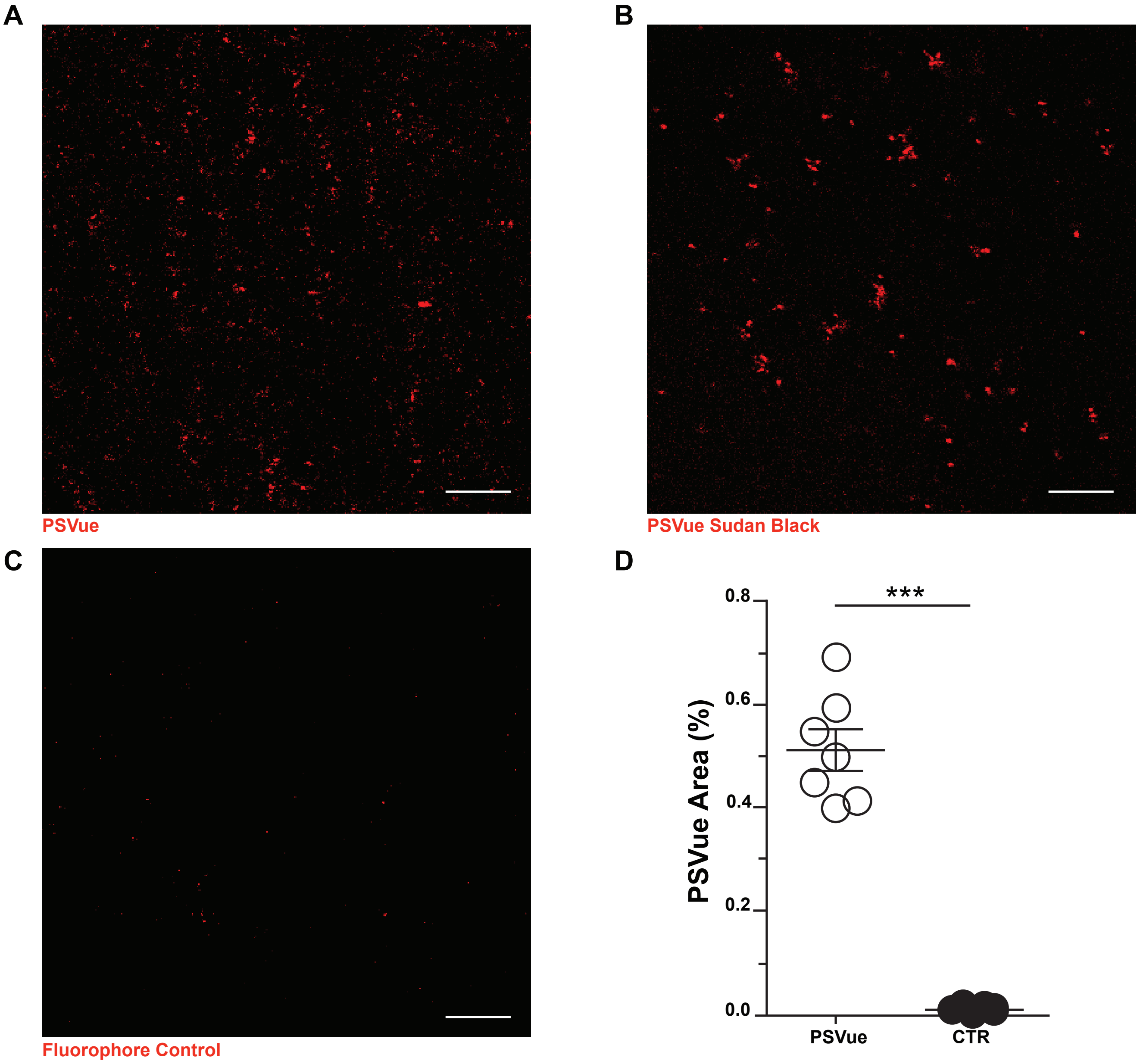


**Supplemental Figure 2: In vivo ePS signal is maintained after aspecific background removal**
(**A-C)** Representative images of CA1 of WT P18 animals sacrificed 3 hours after PSVue injection. (A) Psvue signal was tested by incubating slices with (B) Sudan Black . (C) The specificity of the signal was confirmed by comparing Psvue signal with slices stained with fluorophore only. Scale bar: 5μm.
(**D**) Quantification of PSVue puncta in WT P10 animals compared to unlabeled control. WT PSVue: 0.5149 ± 0.04 vs CTRL: 0.011 ± 0.002; WT PSVue: N=4 animals n=7 fields; CTRL: N= 3 animals n= 6 fields; ***$p<0.001$ Unpaired t-test.
