## Supplemental Figure 3 for "Local externalization of phosphatidylserine mediates developmental synaptic pruning by microglia"

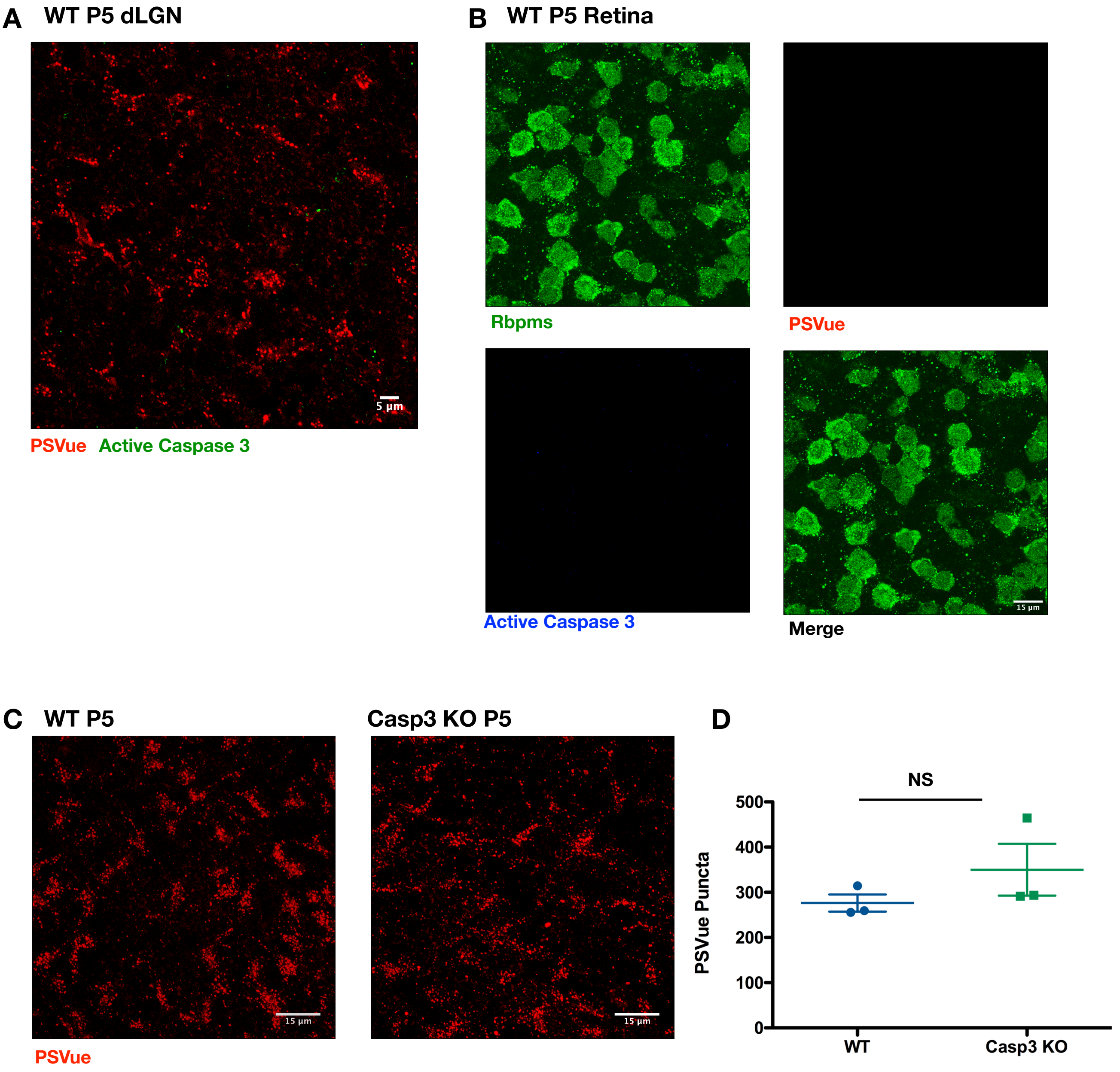


**Supplemental Figure 3:** ***In vivo*** **PS exposure is not downstream of activated caspase 3**
**(A)** Representative max intensity image of the dLGN following injection with PSVue 24 hours prior in WT P5 C57/Bl6 mice. IHC for active caspase3 was performed. Images taken at 63X magnification; scale bar represents 5μm. **(B)** Representative images of retinal whole mounts of WT P5 C57/Bl6 mice following injection with PSVue 24 hours prior. IHC for the pan-RGC marker Rpbms and active caspase3 was performed. Images taken at 63X magnification; scale bar represents 15μm. **(C)** Representative max intensity images of the dLGN following injection with PSVue 24 hours prior in WT or caspase3 knockout (KO) P5 littermates. Images taken at 63X magnification; scale bar represents 15μm. **(D)** Quantification of PSVue in the dLGN of WT or caspase3 KO P5 littermates injected with PSVue 24 hours prior. Data represent the mean per animal ± SEM; N=3; WT: 276.4 ± 19.01 vs. caspase3 KO: 349.9 ± 57.29 p=0.2904 unpaired t-test.
