## Supplemental Figure 4 for "Local externalization of phosphatidylserine mediates developmental synaptic pruning by microglia"

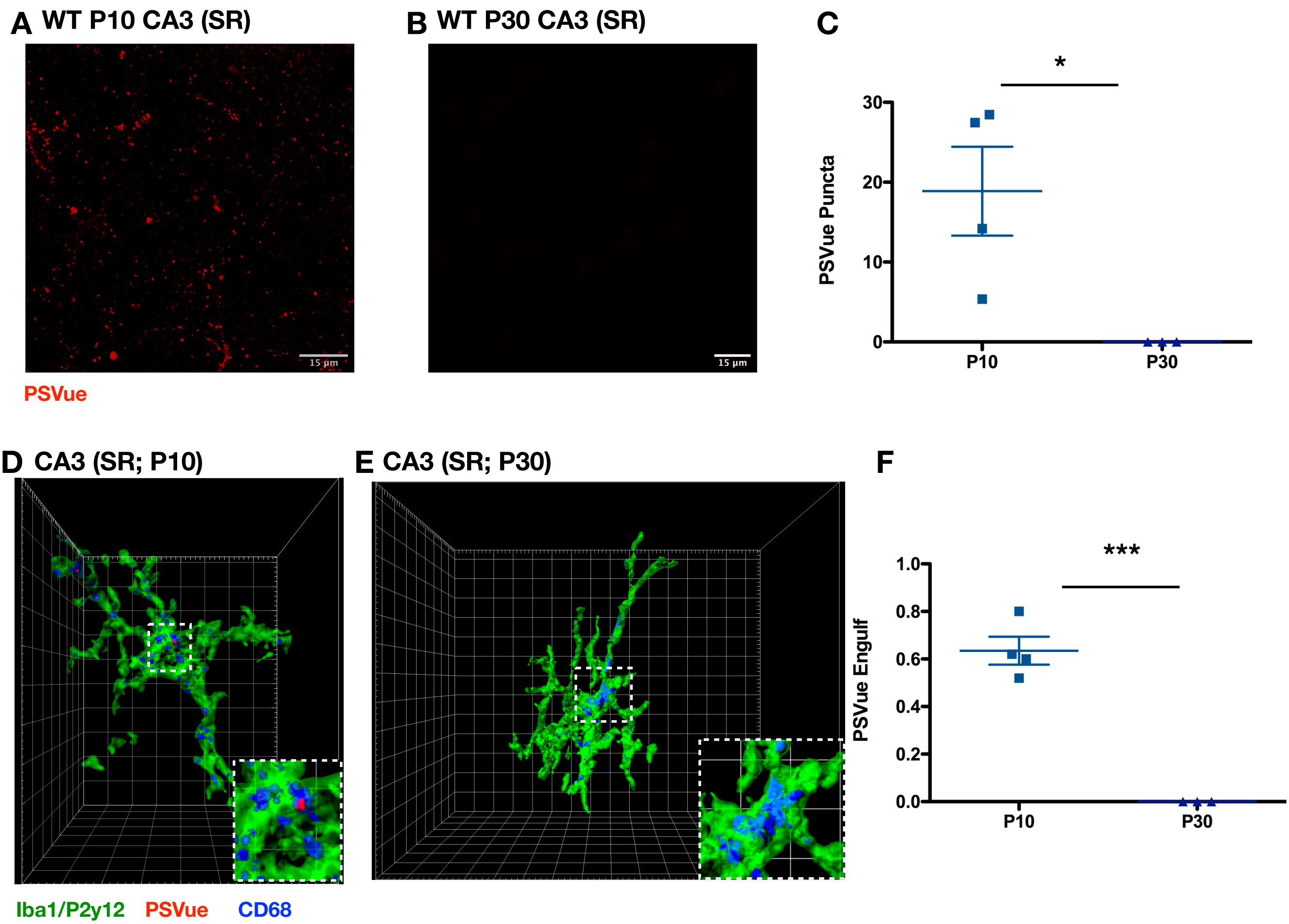


**Supplemental Figure 4: Developmental PS exposure and engulfment in CA3**
**(A, B)** Representative max intensity images of CA3 SR following injection with PSVue 24 hours prior in WT P10 **(A)** and P30 **(B)** C57/Bl6 mice. Images taken at 63X magnification; scale bar represents 15µm.
**(C)** Quantification of PSVue in CA3 SR of WT P10 and P30 C57/Bl6 mice injected with PSVue 24 hours prior. Data represent the mean per animal ± SEM; N=4 (P10), N=3 (P30); P10: 18.08 ± 5.55 vs. P30: $2.34e^{-5}$ ± $1.67e^{-7}$ *p=0.0349 unpaired t-test. **(D, E)** Representative Imaris surface-rendered images of microglia in CA3 SR following injection with PSVue 24 hours prior in WT P10 **(D)** and P30 **(E)** C57/Bl6 mice. Microglia are labeled by IHC with Iba1 and P2y12, and lysosomes are labeled with CD68. Images taken at 63X magnification. **(F)** Quantification of the volume of engulfed PSVue material in CA3 microglia analyzed from WT P10 and P30 C57/Bl6 mice injected with PSVue 24 hours prior. Data represent the mean of 15-20 microglia per animal ± SEM; N=4 (P10), N=3 (P30); P10: 0.635% ± 0.059 vs. P30: $1.35e^{-5}$% ± $1.67e^{-7}$ ***p=0.0003 unpaired t-test.
